# Mutations in ClpC1 or ClpX subunit of the caseinolytic protease confer resistance to natural product ilamycins in mycobacteria

**DOI:** 10.1101/2024.02.24.581832

**Authors:** Yamin Gao, Cuiting Fang, Biao Zhou, H.M. Adnan Hameed, Changli Sun, Xirong Tian, Jing He, Xingli Han, Han Zhang, Jun Li, Jianhua Ju, Xinwen Chen, Nanshan Zhong, Junying Ma, Xiaoli Xiong, Tianyu Zhang

**Author notes:** Corresponding authors: Tianyu Zhang; Xiaoli Xiong; Junying Ma. These authors contributed equally to this work (Yamin Gao, Cuiting Fang).

## Abstract

The mycobacterial caseinolytic protease (Clp) system has been recognized as a promising therapeutic target. In this study, we identified two novel ilamycin analogs, ilamycin E (ILE) and ilamycin F (ILF), both targeting the ClpC1 component of the ClpC1P1P2 proteasome. ILE potently disrupts ClpC1P1P2-mediated proteolysis, leading to delayed bactericidal activity, while ILF also binds ClpC1, albeit with lower affinity. Notably, we discovered a unique mutation in *clpX* and a novel insertion in *clpC1*, both conferring resistance to ILE and ILF in mycobacterium, validated by gene editing. Furthermore, ILE can also inhibit the proteolytic activity of ClpXP1P2 in a manner dependent on the substrate’s tag sequence and adaptor. This first demonstration of *clpX*- and *clpC1*-mediated ilamycin resistance underscores the potential of ilamycins to target multiple components of the Clp protease system, offering a novel dual-target strategy for combating mycobacterial infections.

## Introduction

Tuberculosis (TB) caused by *Mycobactrium tuberculosis* (Mtb) continues to pose a significant global public health threat, with the situation being exacerbated by the emergence of drug-resistant Mtb strains^1,2^. Additionally, infections caused by nontuberculous mycobacteria (NTM), such as *Mycobacterium abscessus* (Mab), are on the rise due to intrinsic resistance to almost all the existing antibiotics. In several countries, the incidence of NTM infections has surpassed that of TB^3,4^. This highlights an urgent need for new drugs with novel mechanisms of action to manage drug-resistant mycobacterial diseases effectively.

The caseinolytic protease (Clp) proteolytic system has recently emerged as a promising therapeutic target in mycobacteria^5–8^. This system is pivotal not only for maintaining intracellular protein quality but also for modulating responses to environmental stressors and contributing to the pathogenicity of virulent strains^6,9^. It comprises multi-subunit protein complexes that facilitate intracellular protein degradation. In mycobacteria, the core complex of the hetero-tetradecamer Clp protease assumes a barrel-shaped structure, constructed from two rings, each composed of seven ClpP1 and seven ClpP2 peptidase subunits. The entry of substrates into the ClpP1P2 complex is stringently regulated and reliant on ATP- dependent AAA+ unfoldases (ClpX or ClpC1), which act as adaptors to facilitate substrate entry into the protease chamber through ATP hydrolysis^8, 10, 11^. Both the Clp protease genes and the ATPase adaptor genes *clpX* and *clpC1* have been found essential for mycobacterial survival^8,12^.

Many non-ribosomal natural cyclic peptides, such as ilamycins (also known as rufomycins), cyclomarin A (CYMA), and ecumicin (ECU), have shown antitubercular activities by targeting the ClpC1 component of the ClpC1P1P2 proteasome^13^. However, aside from the mutations identified in the *clpC1* gene from spontaneous resistant mutants, robust genetic evidence is lacking. Although these compounds all disrupt protein metabolism through the Clp system, their mechanisms of action (MOAs) differ. For example, rufomycin □ (RUFI) inhibits the proteolytic activity of the ClpP1P2 complexes by binding to ClpC1, interfering with its binding to ClpP1P2, while CYMA stimulates ClpC1’s ATPase and promotes proteolysis by the ClpC1P1P2 proteasome^14,15^. On the other hand, ECU stimulates the ATPase activity of ClpC1 but inhibit the proteolysis by ClpC1ClpP1P2^16^. To the best of our knowledge, there are no reported anti-Mtb compounds targeting ClpX.

We previously reported that ilamycin E (ILE) and ilamycin F (ILF), synthesized by *Streptomyces atratus* SCSIO ZH16 isolated from the South China Sea, exhibited activities against Mtb^17^. Particularly, ILE demonstrated superior activity against Mtb compared to other ilamycin components and the aforementioned compounds^13,17,18^. However, their MOAs, activities against NTM and other non-mycobacterial pathogenic bacteria, and potential drug interactions remain to be elucidated. In this study, we discovered that ilamycins target ClpC1 and inhibit the proteolytic activity of the ClpC1P1P2 protease. Additionally, ILE can also impair the proteolytic activity of the ClpXP1P2 protease depending on the tag sequences of the substrate and adaptor-mediated recognition. These findings highlight the potential of Clp complexes as viable therapeutic targets, particularly when employing a dual-targeting approach.

## Results

### ILE and ILF exhibit selective and potent antimycobacterial activities *in vitro* and *ex vivo*

Mtb and 3 different NTM (Mab, *Mycobacterium marinum* (Mmr), *Mycobacterium smegmatis* (Msm) were selected to investigate the activities of ILE and ILF. These mycobacteria were chosen due to their clinical relevance and growth characteristics. Mab, a rapidly growing mycobacterium, is frequently associated with severe human infections and is known for its intrinsic antibiotic-resistant nature^19,20^. Mmr, a slowly growing mycobacterium, is closely related to Mtb but less virulent and serves as a model organism for Mtb studies^21,22^. Msm is also utilized widely as a model organism due to its rapid growth, non-virulence and the availability of genetic tools^22^.

ILE and ILF demonstrated significant inhibitory activities against autoluminescent Mtb H_37_Ra (AlRa)^23^, Msm (AlMsm)^23^ and Mmr (AlMmr)^24^ with varying potencies (Table 1). Notably, ILE exhibited potent activity against autoluminescent Mab (AlMab)^25^ at a concentration of 1 µg mL^-1^, whereas ILF only showed weak activity at 20 µg mL^-1^, though they both showed highly bactericidal activities against Mtb. This disparity in efficacy may stem from ILE and ILF structural differences and the unique cell wall permeabilities of each mycobacterial species^26,27^. Compared to RIF, ILE and ILF exhibited delayed antimicrobial activities, initially displaying minor bacteriostatic effects on AlRa, with bactericidal activity emerging after 6 days of incubation (Fig. 1). Similar killing dynamics were observed for AlMsm, AlMab, and AlMmr (Supplementary Fig. 1). Consistent bactericidal activity of ILE was also confirmed in autoluminescent H_37_Rv (AlRv) (Supplementary Fig. 2), addressing concerns regarding the slower growth of H_37_Ra^28^. The classical colony-forming unit (CFU) enumeration were also performed to evaluate the antimicrobial activities of ILE and ILF. Consistent with the luminescence inhibition data, ILE and ILF exhibited concentration- and time-dependent bactericidal activities against AlRa, as demonstrated by the substantial reduction in CFU counts after 15 days of incubation (Fig. 1). Specifically, based on CFU enumeration in AlRa, the minimal bactericidal concentration (MBC) was determined to be 0.05 μg mL^-1^ for ILE and 1.25 μg mL^-1^ for ILF.

**Fig 1.**
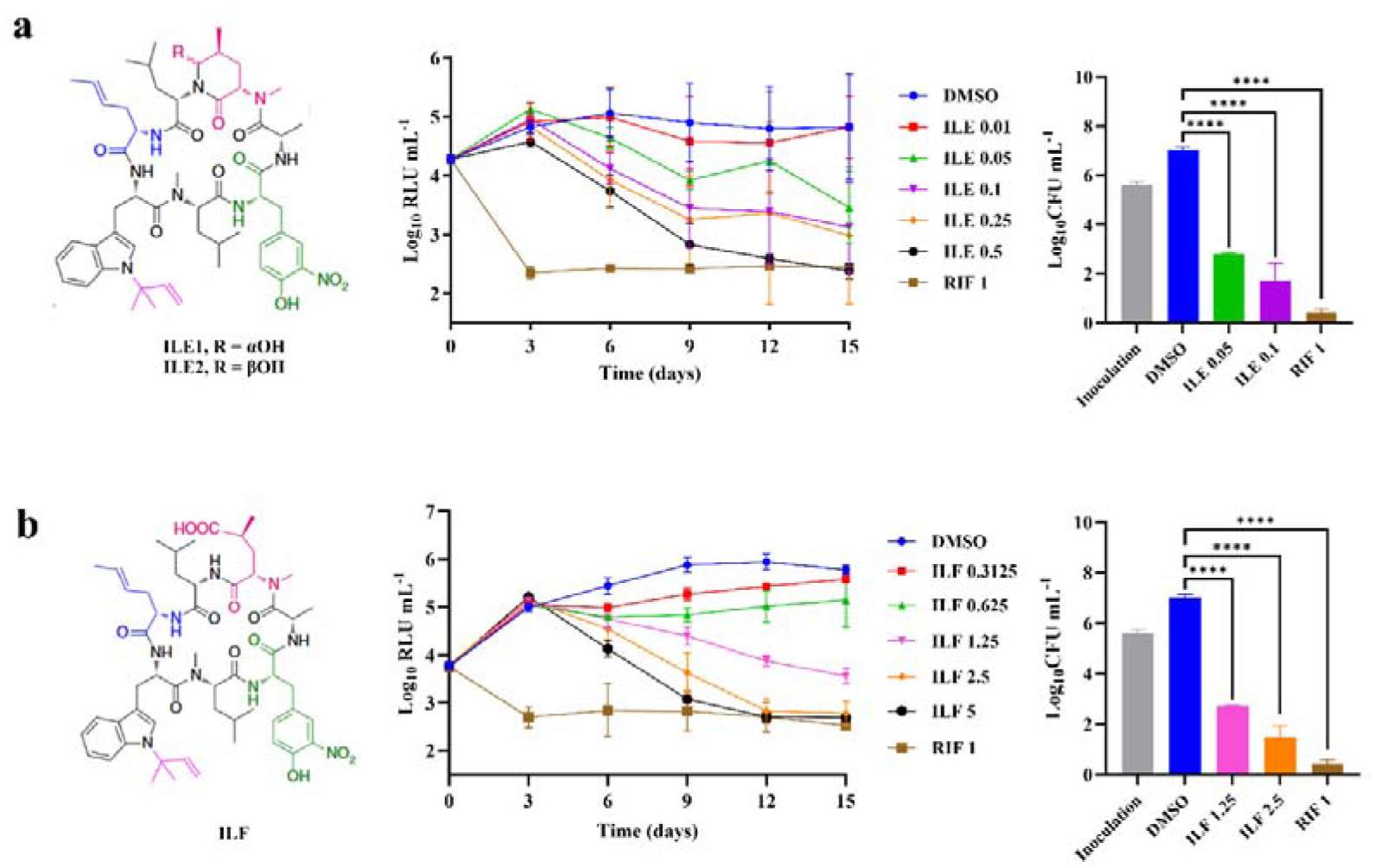
LE and ILF exhibit potent activity against AlRa. Chemical structures, time-kill curves based on relative light units (RLUs), and CFU counts at day 15 against AlRa of ILE **(a)** and ILF **(b)**. DMSO, solvent control; RIF, positive control. Numbers following the drug names indicate the working concentrations (µg mL^-1^). The MIC_lux_ was defined as the lowest concentration that can inhibit > 90% RLUs compared with that from the untreated controls. RLU and CFU data were obtained from independent experiments. Statistical significance was determined using one-way ANOVA. Data are presented as mean ± SD from three biological replicates (n = 3). ****, *P* < 0.0001.

**Table 1.** Antibacterial spectrum of ILE and ILF.

| Strains or drug combinations | ILE MIC ($\mu\text{g mL}^{-1}$) | ILF MIC ($\mu\text{g mL}^{-1}$) | FICI |
| --- | --- | --- | --- |
| AlRa | 0.05<sup>†</sup> | 1.25<sup>†</sup> | - |
| AlRv | 0.05<sup>†</sup> | ND | - |
| AlMsm | 0.0625<sup>†</sup> | 2.5<sup>†</sup> | - |
| AlMab | 1<sup>†</sup> | 20<sup>†</sup> | - |
| AlMmr | 0.08<sup>†</sup> | 0.625<sup>†</sup> | - |
| H<sub>37</sub>Rv | 0.025<sup>‡</sup> | 2.5<sup>‡</sup> | - |
| K11<sup>*</sup> | 0.025-0.05<sup>‡</sup> | ND | - |
| K12<sup>*</sup> | 0.025<sup>‡</sup> | ND | - |
| K13<sup>*</sup> | 0.01<sup>‡</sup> | ND | - |
| K14<sup>*</sup> | 0.01<sup>‡</sup> | ND | - |
| K16<sup>*</sup> | 0.05<sup>‡</sup> | ND | - |
| K17<sup>*</sup> | 0.01<sup>‡</sup> | ND | - |
| K18<sup>*</sup> | 0.025-0.05<sup>‡</sup> | ND | - |
| K19<sup>*</sup> | 0.01<sup>‡</sup> | ND | - |
| K20<sup>*</sup> | 0.05<sup>‡</sup> | ND | - |
| *Staphylococcus aureus* | > 128 | > 128 | - |
| *Acinetobacter baumannii* | > 128 | > 128 | - |
| *Enterococcus faecium* | > 128 | > 128 | - |
| *Klebsiella pneumoniae* | > 128 | > 128 | - |
| *Pseudomonas aeruginosa* | > 128 | > 128 | - |
| ILE + RIF | - | - | 0.3125<sup>†</sup> |
| ILE + INH | - | - | 0.625<sup>†</sup> |
| ILE + EMB | - | - | 0.5625<sup>†</sup> |
| ILE + STR | - | - | 1<sup>†</sup> |
| ILE + CFZ | - | - | 1<sup>†</sup> |
| ILE + BDQ | - | - | 0.62<sup>†</sup> |
| ILE + TB47 | - | - | 0.3125<sup>†</sup> |
| ILF + RIF | - | - | 1.0625<sup>†</sup> |
| ILF + INH | - | - | 0.3125<sup>†</sup> |
| ILF + EMB | - | - | 1<sup>†</sup> |
| ILF + STR | - | - | 1<sup>†</sup> |
| ILF + CFZ | - | - | 0.63<sup>†</sup> |
| ILF + BDQ | - | - | 1<sup>†</sup> |
| ILF + TB47 | - | - | 0.3125<sup>†</sup> |

\*Clinical resistant Mtb; <sup>†</sup>MICs obtained by detection of relative light units; <sup>‡</sup>MICs obtained by the microplate alamar blue assay; -Not applicable; ND, not detected ; FICI, fractional inhibitory concentration index.

The minimal inhibitory concentrations (MICs) of ILE against clinically isolated drug-resistant Mtb strains ranged from 0.01 to 0.05 µg mL^-1^, which aligns with the laboratory standard Mtb H_37_Rv strain (Table 1). Synergistic effects were observed when combining ILE with rifampicin (RIF) or the novel anti-TB drug TB47 or combining ILF with isoniazid (INH) or TB47^29^. Partial synergy was noted when ILE was paired with INH, ethambutol (EMB), clofazimine (CFZ), or bedaquiline (BDQ), and when ILF was combined with CFZ. Additive effects were observed when ILE was combined with streptomycin (STR), or ILF combined with EMB, STR or BDQ. No antagonism was detected between any of the selected drugs and either ILE or ILF. In mouse RAW264.7 macrophages, ILE effectively inhibited AlRv^30^ with an MIC of 0.2 μg mL^-1^, while ILF showed activity at 5 μg mL^-1^. Cytotoxicity testing of ILE in RAW264.7 cells revealed a 50% toxic concentration (TC_50_) of 28.24□μg□mL^-1^, suggesting a favorable selectivity index (TC_50_/MIC = 141.2) relative to its anti-Mtb activity. However, neither ILE nor ILF exhibited activity against common non- mycobacterial clinical isolates, with MICs exceeding 128 µg mL^-1^ (Table 1).

### Mutations in *clpC1* and *clpX* were identified in ILE/ILF-resistant mutants

To identify target(s) of ILE and ILF, spontaneous mutants of different mycobacterium species were screened. The spontaneous mutation rate for producing ILE-resistant Mtb was approximately 2.5 × 10^-9^, which is similar to that of RUFI and lower than that of ECU and lassomycin, as well as RIF^15,16,18,31^.

Whole genome sequencing (WGS) of 7 ILE-resistant Mtb strains from 4 independent screens revealed no mutations in ClpC1 or its flanking regions, but 5 strains carried a ClpX P30H mutation. Further Sanger sequencing of 17 ILE-resistant Mtb strains identified ClpX (P30H) mutation in 8 strains (47%) and ClpC1 (F80V/C/I) mutations in 11 strains (61%), with 2 strains harboring both mutations (ClpX P30H and ClpC1 F80V/I) (Table 2). Similarly, WGS of 8 ILE-resistant Mab strains revealed 4 strains with ClpC1 mutations (Q17R, F2L, H77D), one with ClpX P30H, and three without mutations in either gene. Sanger sequencing of the *clpX* and *clpC1* genes of other ILE-resistant Mab strains revealed that there were also ClpC1 (M1-FERF-F2; L88R) mutations in Mab strains (Table 2). Besides, a small subset of ILE-resistant Mtb or Mab strains did not carry mutations in neither *clpX* nor *clpC1*, suggesting the potential involvement of alternative resistance mechanisms. In these strains, WGS identified sporadic mutations in genes unrelated to the Clp proteases, such as *Rv1572*, *MAB_2872c* (*hisS*), or *MAB_2670c* (*hisD*) (Supplementary Table 1 and Supplementary Table 2), though their relevance to resistance remains to be validated.

**Table 2.** Summary of mutations of ClpC1 and ClpX in ILE- and ILF-resistant mutants.

| Strains No.<sup>a</sup> | MICs (μg mL<sup>-1</sup>) |  | Mutation |  |  |  |
| --- | --- | --- | --- | --- | --- | --- |
|  | ILE | ILF | ClpC1 | *clpC1* | ClpX | *clpX* |
| Mtb<sup>R1*</sup> | 0.625 | - | - | - | P30H | CCC→C<u>A</u>C |
| Mtb<sup>R2*</sup> | 20 | - | F80I | <u>T</u>TT→<u>A</u>TT | P30H | CCC→C<u>A</u>C |
| Mtb<sup>R3*</sup> | 20 | - | F80V | <u>T</u>TT→<u>G</u>TT | P30H | CCC→C<u>A</u>C |
| Mtb<sup>R4*</sup> | 0.1 | - | F80I | <u>T</u>TT→<u>A</u>TT | - | - |
| Mtb<sup>R5*</sup> | 0.1 | - | F80V | <u>T</u>TT→<u>G</u>TT | - | - |
| Mtb<sup>R6*</sup> | 0.1 | - | F80L | <u>T</u>TT→<u>C</u>TT | - | - |
| Mtb<sup>R7*</sup> | 20 | - | F80C | <u>T</u>TT→T<u>G</u>T | - | - |
| Mab<sup>R1*</sup> | > 128 | > 128 | - | - | P30H | CCC→C<u>A</u>C |
| Mab<sup>R2*</sup> | 4 | > 128 | F2L | <u>T</u>T<u>C</u>→T<u>C</u><u>C</u> | - | - |
| Mab<sup>R3*</sup> | 4 | > 128 | M1-FERF-F2 | Ins: | - | - |
|  |  |  |  | TTCGAGAG<br>ATTC |  |  |
| Mab<sup>R4*</sup> | 4 | > 128 | Q17R | <u>C</u>AA→<u>C</u>GA | - | - |
| Mab<sup>R5*</sup> | > 128 | > 128 | H77D | <u>C</u>AC→<u>G</u>AC | - | - |
| Mab<sup>R6*</sup> | > 128 | > 128 | L88R | <u>C</u>T<u>G</u>→<u>C</u><u>G</u><u>G</u> | - | - |
| Mmr<sup>R1*</sup> | 1 | 4 | F2L | <u>T</u>T<u>C</u>→T<u>C</u><u>C</u> | - | - |
| Mmr<sup>R2*</sup> | 10 | 40 | H77D | <u>C</u>AC→<u>G</u>AC | - | - |
| Mmr<sup>R3*</sup> | 10 | 40 | L88R | <u>C</u>T<u>C</u>→<u>C</u><u>G</u><u>C</u> | - | - |
| Msm<sup>R1†</sup> | 0.5 | 20 | F2S | <u>T</u>TT→T<u>C</u>T | - | - |
| Msm<sup>R2†</sup> | 0.25 | 10 | F2C | <u>T</u>TT→T<u>G</u>T | - | - |
| Msm<sup>R3†</sup> | 1 | 20 | M1-L-F2 | Ins: GTT | - | - |
| Msm<sup>R4†</sup> | 1 | 10 | F5I | <u>T</u>TT→<u>A</u>TT | - | - |
| Msm<sup>R5†</sup> | 0.5 | 40 | F80S | <u>T</u>T<u>C</u>→T<u>C</u><u>C</u> | - | - |
| Msm<sup>R6†</sup> | 0.25 | 10 | L88P | <u>C</u>T<u>C</u>→<u>C</u><u>C</u><u>C</u> | - | - |
| Msm<sup>R7†</sup> | 0.25 | 20 | L88R | <u>C</u>T<u>C</u>→<u>C</u><u>G</u><u>C</u> | - | - |
| Msm<sup>R8†</sup> | 0.25 | 10 | F2S; I197N | <u>T</u>TT→T<u>C</u>T; | - | - |
|  |  |  |  | <u>A</u>T<u>C</u>→<u>A</u><u>A</u><u>C</u> |  |  |

<sup>a</sup>Resistant strains were renumbered; <sup>\*</sup>Resistant mutants selected from ILE-containing plates; <sup>†</sup>Resistant mutants selected from ILF-containing plates; Ins, insertion; Underlines indicate mutated bases; - showed no mutation.

However, all examined ILE-resistant Mmr strains and ILF-resistant Msm strains carried mutations exclusively in ClpC1 without any mutations in ClpX (Table 2). Cross-resistance between ILE and ILF was observed in all ILE- or ILF-resistant NTM mutants. These findings suggest that ILE and ILF may both target the ClpX and ClpC1, especially in Mtb and Mab.

### Docking predicts ILE and ILF bind the N-terminal domain (NTD) of ClpC1 and ClpX

To investigate the interactions of ILE and ILF with ClpC1 and ClpX, we performed molecular docking simulations. Since all mutation sites of ILE and ILF- resistant bacteria are located in ClpC1-NTD (residues 1–145) or ClpX-NTD (residues 1–112), we employed AutoDock to dock ILE into the reported structure of the MtbClpC1-NTD (Fig. 2a)^32–34^ and the AlphaFold2-predicted structure of the MtbClpX-NTD (Fig. 2b), respectively. The docking results indicated that ILE binds to MtbClpC1-NTD and MtbClpX-NTD with predicted binding energies of –7.9 kcal mol^-1^ and –6.7 kcal mol^-1^, and corresponding root mean square deviations of 3.5 Å and 2.1 Å, respectively, confirming acceptable docking accuracy. Key residues involved in ILE binding include the known mutation sites Q17, H77, F80, and L88 in MtbClpC1-NTD, and P30 in MtbClpX-NTD (Table 2). Given the structural similarity between ILE and ILF, along with the observed cross-resistance, it is likely that ILF interacts with ClpC1 and ClpX at similar binding sites. Notably, H77 and F80 are also critical for CYMA and RUFI binding, while RUFI additionally involves M1, F2, V13, and E89, and ECU interacts with L92 and L96, indicating a partially conserved binding interface across cyclopeptides^32^.

**Fig 2.**
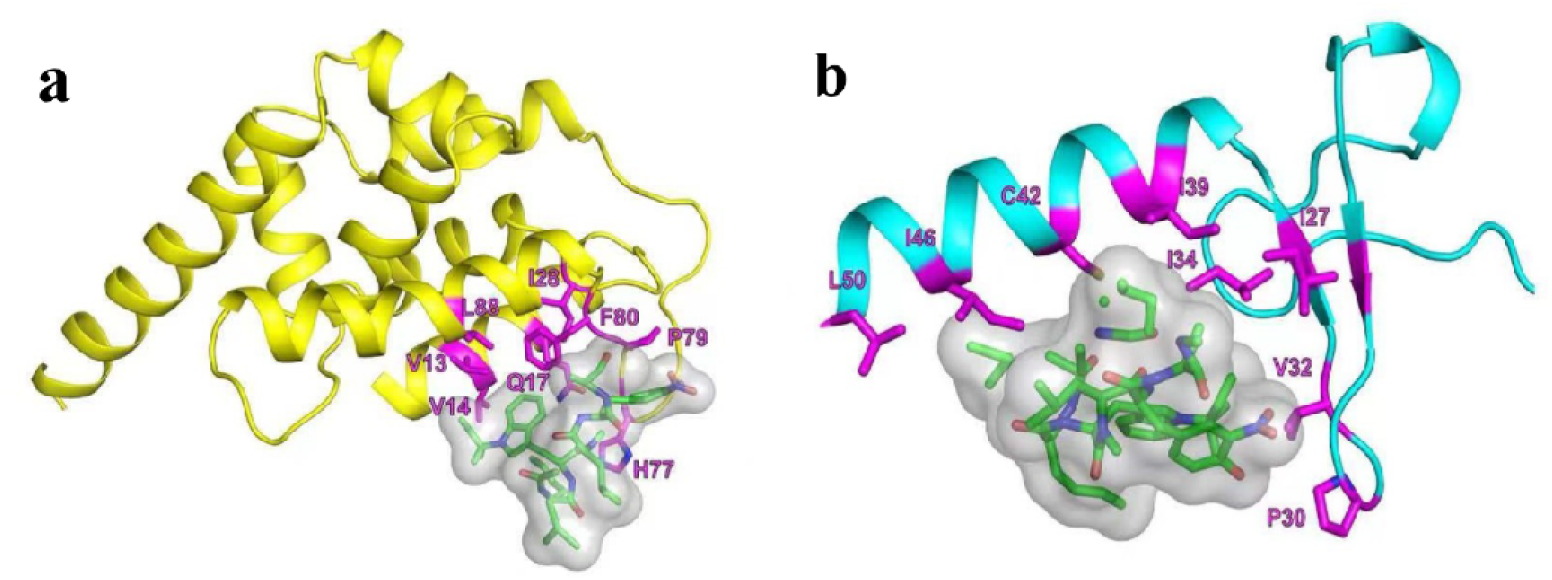
The graphical representation of crucial residues in the interaction between ILE and MtbClpC1-NTD as well as MtbClpX-NTD. **a** Crucial residues in the interaction between ILE and MtbClpC1-NTD. **b** Crucial residues in the interaction between ILE and MtbClpX-NTD. MtbClpC1-NTD, N-terminal domain of MtbClpC1 (residues 1–145); MtbClpX-NTD, N-terminal domain of MtbClpX (residues 1–112).

### Mutations in *clpC1* and *clpX* confer resistance to ILE and ILF in mycobacteria

To validate the significance of ClpC1 and ClpX in relation to ILE and ILF, we initially constructed a series of overexpression strains. The wild-type (wt) genes *clpX*^wt^ and *clpC1*^wt^, along with their mutant (mt) variants *clpX*^mt^ and *clpC1*^mt^, were inserted into two extrachromosomal plasmids, p60A or pMVA (Supplementary Fig. 3)^35^. These genes were under the robust *hsp60* promoter to facilitate overexpression in AlMab, AlMsm, and AlMmr^23–25^. Among these strains, only the AlMmr strain overexpressing ClpC1^L88R^ demonstrated resistance to both ILE and ILF when compared to the parent strain, exhibiting an 8-fold increase in MICs (Supplementary Table 3). However, the overexpression of ClpC1^wt/mt^ or ClpX^wt/mt^, as well as the concurrent overexpression of genes encoding the ClpC1P1P2 or ClpXP1P2 proteases in AlMab and AlMsm, did not alter their susceptibility to ILE or ILF. These findings are consistent with previous observations, suggesting that the overexpression of all mutated *clpC1* does not necessarily confer resistance to ClpC1-targeting compounds, and that efficacy varies across different mycobacterial species^36^.

In order to further determine the roles of mutations in *clpC1* and *clpX* in ILE and ILF resistance directly, we aimed to obtain gene-edited strains. Given the slower growth rate of Mtb compared to rapidly-growing mycobacteria such as Msm and Mab, we initially utilized these two NTM^37^. Utilizing the newly developed CRISPR/Cpf1- mediated gene editing techniques, we introduced specific *clpC1* and *clpX* mutations into Mab and Msm, resulting in two Mab mutants (ClpC1^wt^ to ClpC1^F2L^; ClpX^wt^ to ClpX^P30H^) and three Msm mutants (ClpC1^wt^ to ClpC1^F2C^, ClpC1^wt^ to ClpC1^M1-L-F2^ and ClpC1^wt^ to ClpC1^I197N^)^38,39^. These genetically modified strains displayed varying levels of resistance as summarized in Table 3. Notably, the ClpX^P30H^ mutation in Mab conferred the highest level of resistance, with MICs exceeding 128 µg mL^-1^ for both ILE and ILF. The ClpC1^F2L^ mutation in Mab led to a more modest increase in resistance, elevating the MIC for ILE by eight times relative to the parent strain. The mutations ClpC1^F2C^ and ClpC1^M1-L-F2^ in Msm both resulted in a greater than 4-fold increase in the MICs of ILE and ILF, compared to the parent strain. A comparative analysis of spontaneous mutants and gene-edited strains revealed that mutations at the F2 of ClpC1 in both Mab and Msm confer relatively low-level resistance to ILE and ILF, whereas the P30H mutation in Mab ClpX is associated with high-level resistance, underscoring its crucial role. It is noteworthy that the lack of impact of the Msm ClpC1^I197N^ mutation on ILE and ILF resistance may be attributed to its location outside ClpC1-NTD, suggesting that resistance is primarily driven by alterations within the NTD. Furthermore, the spontaneous mutant strain harboring *clpC1*^I197N^ also possessed *clpC1*^F2S^ (Table 2) which may be the principal factor contributing to ILE and ILF resistance.

**Table 3.** MICs of ILE and ILF against gene edited strains and corresponding spontaneous mutants.

| Strains | Amino acid change | MICs ($\mu\text{g mL}^{-1}$) / (Fold change) | |
| --- | --- | --- | --- |
|  |  | ILE | ILF |
| Mab<sup>wt</sup> | / | 1 | 20 |
| Mab<sup>R2*</sup> | ClpC1<sup>wt</sup> to ClpC1<sup>F2L</sup> | 4 / (4) | > 128 / (> 6) |
| Mab-C1<sup>F2L</sup> | ClpC1<sup>wt</sup> to ClpC1<sup>F2L</sup> | 8 / (8) | > 128 / (> 6) |
| Mab<sup>R1*</sup> | ClpX<sup>wt</sup> to ClpX<sup>P30H</sup> | > 128 / (> 128) | > 128 / (> 6) |
| Mab-X<sup>P30H</sup> | ClpX<sup>wt</sup> to ClpX<sup>P30H</sup> | > 128 / (> 128) | > 128 / (> 6) |
| Msm<sup>wt</sup> | / | 0.0625 | 2.5 |
| Msm<sup>R2†</sup> | ClpC1<sup>wt</sup> to ClpC1<sup>F2C</sup> | 0.25 / (4) | 10 / (4) |
| Msm-C1<sup>F2C</sup> | ClpC1<sup>wt</sup> to ClpC1<sup>F2C</sup> | 0.5 / (8) | 20 / (8) |
| Msm<sup>R3†</sup> | ClpC1<sup>wt</sup> to ClpC1<sup>M1-L-F2</sup> | 1 / (16) | 20 / (8) |
| Msm-C1<sup>M1-L-F2</sup> | ClpC1<sup>wt</sup> to ClpC1<sup>M1-L-F2</sup> | 0.25 / (4) | 16 / (6.4) |
| Msm<sup>R8†</sup> | ClpC1<sup>wt</sup> to ClpC1<sup>F2S</sup> | 0.25 / (4) | 10 / (4) |
|  | ClpC1<sup>wt</sup> to ClpC1<sup>I197N</sup> |  |  |
| Msm-C1<sup>I197N</sup> | ClpC1<sup>wt</sup> to ClpC1<sup>I197N</sup> | 0.0625 / (1) | 2.5 / (1) |

\*Resistant mutants selected from ILE-containing plates; †Resistant mutants selected from ILF-containing plates; -C1 or -X means ClpC1 or ClpX was edited.

### ILE and ILF treatment alter Msm cell morphology and increases sensitivity in ***clpC1* and *clpX* knockdown strains**

To elucidate the bactericidal mechanisms of ILE and ILF, we detected the effect of ILE or ILF treatment on the morphology of Msm cells. Msm was cultured with 0.125 µg mL^-1^ of ILE or 5 µg mL^-1^ of ILF for 15 hours (h). As a control, Msm was also exposed to 1 µg mL^-1^ clarithromycin (CLR), a macrolide antibiotic that inhibits bacterial protein synthesis by specifically targeting the 50S subunit of the ribosome. The lengths of CLR-treated cells showed only a modest increase, from 2.07 ± 0.48 μm to 2.64 ± 0.80 μm (Fig. 3a and Fig. 3b). Notably, the cell lengths of Msm increased to 5.95 ± 2.16 μm and 9.41 ± 2.40 μm following treatment with ILE and ILF, respectively (Fig. 3c and Fig. 3d). In comparison to untreated and CLR-treated Msm, ILE and ILF both induced a pronounced inhibition of cell division, as evidenced by the formation of division septa (Fig. 3c and Fig. 3d). This observation is consistent with the phenotype observed in Msm upon *clpX* gene silencing, and the established role of ClpX in regulating FtsZ assembly and Z-ring formation which are crucial processes for cell division in Mtb^40,41^.

**Fig 3.**
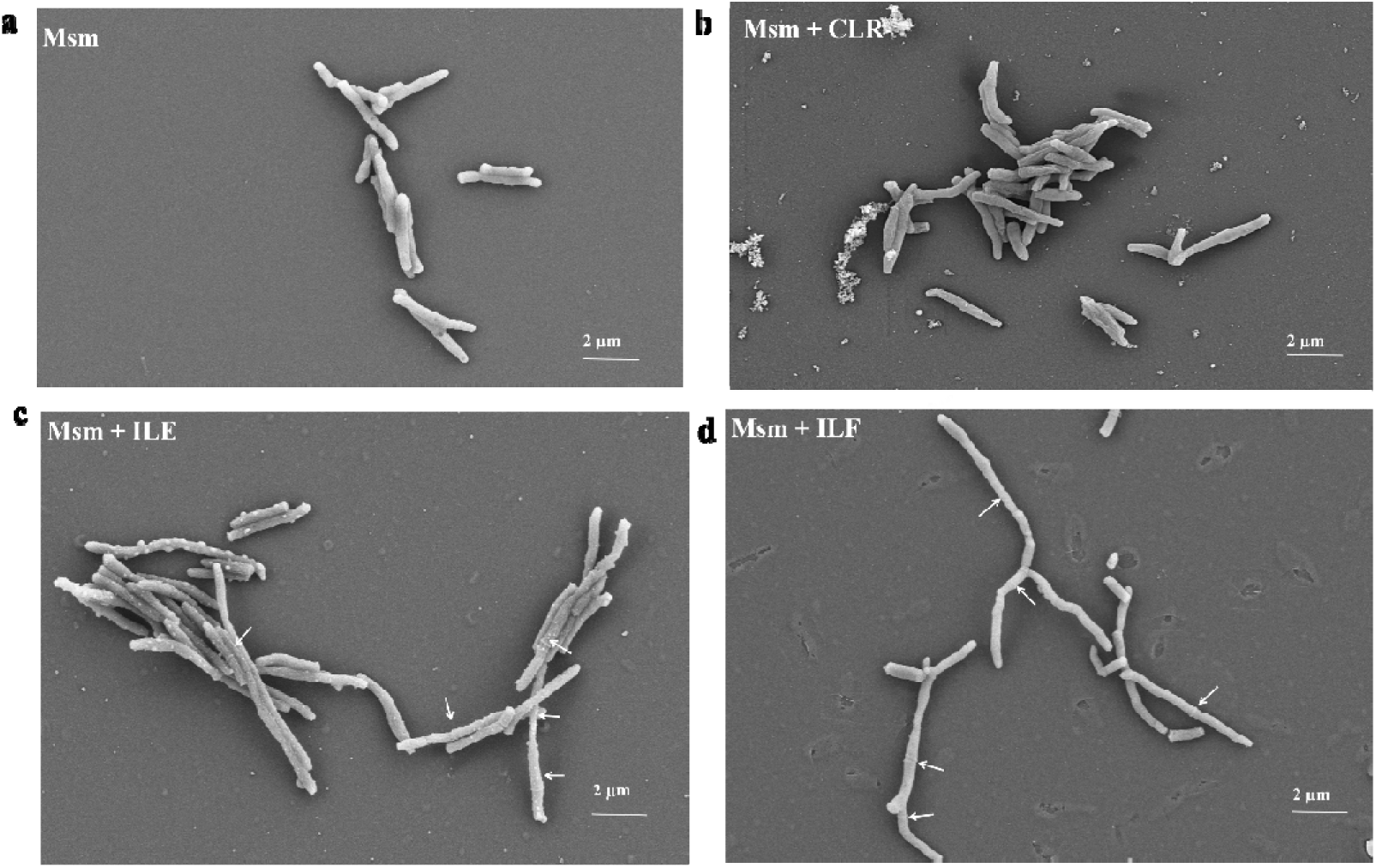
The cell morphology of Msm strains before or after treatment. **a** The cell morphology of Msm without any drug treatment. **b** The cell morphology of Msm treated with a concentration of 1 µg mL^-1^ CLR. **c** The cell morphology of Msm treated with a concentration of 0.125 µg mL^-1^ ILE. **d** The cell morphology of Msm treated with a concentration of 5 µg mL^-1^ ILF. The white arrows highlight the division septa.

Correspondingly, we performed knockdown of *clpX* and *clpC1* in Mab and Msm^42^. The knockdown of *clpX* or *clpC1* in Mab and Msm strains resulted in heightened sensitivity to ILE and ILF (Fig. 4a and Fig. 4b). Upon induction with anhydrotetracycline (aTc) in the absence of ILE or ILF, growth inhibition was observed in both the *clpX* or *clpC1* knockdown strains of Msm and Mab, highlighting the crucial role of these proteins in cellular proliferation. Additionally, we examined ILE’s intracellular activities against *clpC1* and *clpX* knockdown strains of Mab in murine RAW264.7 macrophages (Fig. 4c). Upon infection and drug treatment, ILE exhibited potent intracellular activity, with significantly enhanced efficacy against *clpX* and *clpC1* knockdown strains compared to the vector control. Conclusively, ILE and ILF may disrupt essential cellular processes regulated by ClpX and ClpC1, leading to bactericidal effects.

**Fig 4.**
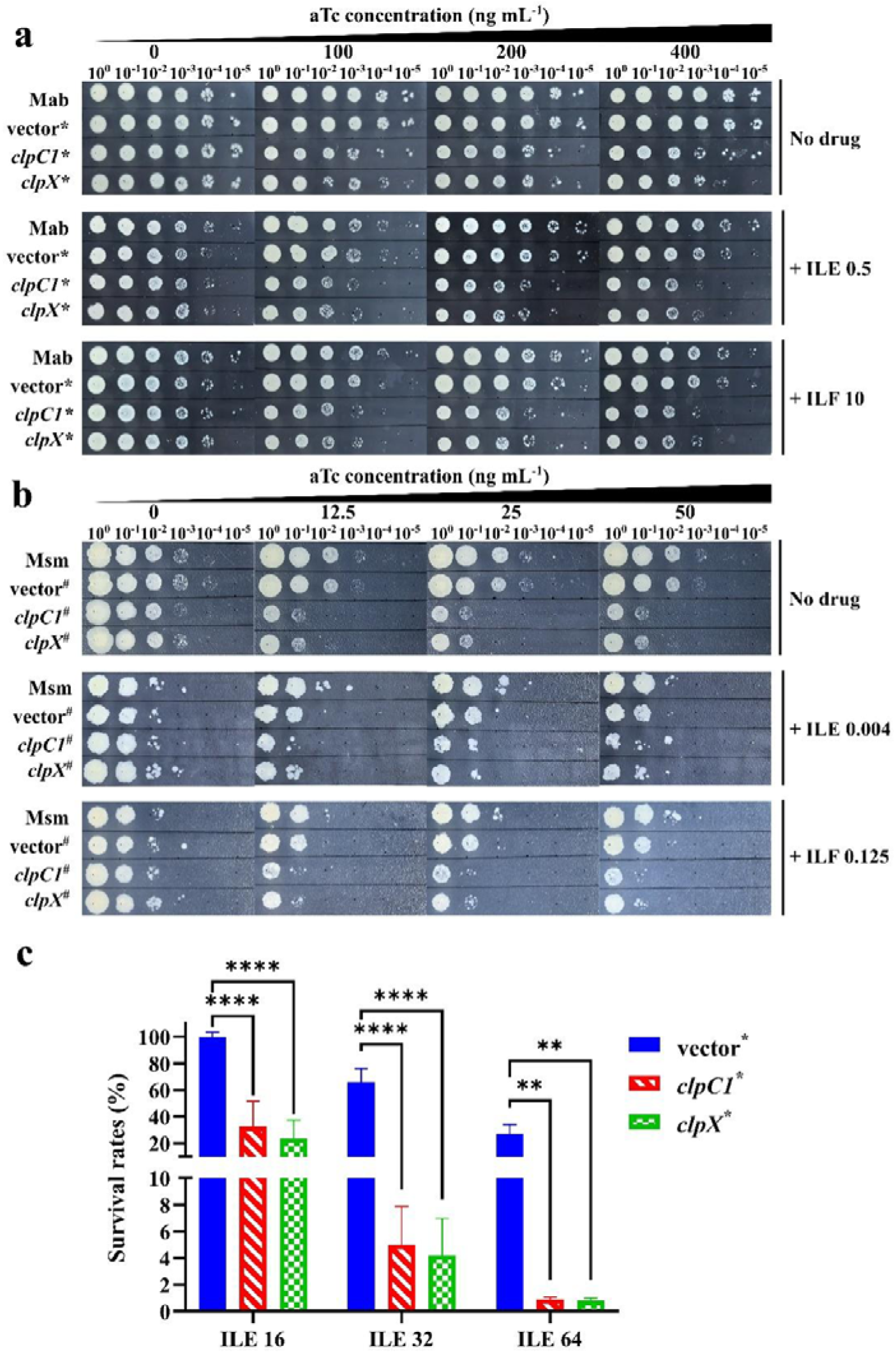
Impact of *clpX* or *clpC1* gene suppression on the susceptibility of Mab and Msm to ILE or ILF. **a** *In vitro* susceptibility of Mab to ILE or ILF upon aTc-induced gene suppression. **b** *In vitro* susceptibility of Msm to ILE or ILF upon aTc-induced gene suppression. The concentrations of ILE and ILF are in μg mL^-1^. *, Mab strains contained varying plasmids: vector, a vector control with the empty plasmid pLJR962; *clpC1*, pLJR962-clpC1 for inducible inhibition of the *clpC1* gene; *clpX*, pLJR962-clpX for inducible inhibition of the *clpX* gene. ^#^, Similarly, Msm strains were differentiated by the plasmids they harbored as that in Mab. **c** Intracellular susceptibility of Mab to ILE upon gene suppression in RAW264.7 macrophages. Survival rates were calculated as (CFU of ILE-treated group / CFU of untreated group) × 100%. Numbers following the drug names indicate the working concentrations (µg mL^-1^). Statistical significance was determined by two-way ANOVA followed by Tukey’s multiple comparisons test. Data are presented as mean ± SD from three biological replicates (n = 3). **, *P* < 0.01; ****, *P* < 0.0001.

### ILE and ILF selectively binds to ClpC1-NTD and is disrupted by resistance mutations

Since all mutations occurred within ClpC1-NTD or ClpX-NTD across Mtb, Mab, Msm, and Mmr, and their ClpC1-NTD or the first 64 amino acids of the ClpX-NTD sequences are identical (Supplementary Fig. 4 and Supplementary Fig. 5), we specifically expressed and purified the MtbClpC1-NTD^wt/mt^ and MtbClpX-NTD^wt/mt^. Our goal was to investigate whether there is any interaction between these proteins and ILE or ILF. The interactions between the MtbClpC1-NTD or MtbClpX-NTD with ILE or ILF were examined by differential scanning fluorimetry (DSF)^43^. We discovered that ILE increased the melting temperature (Tm) of MtbClpC1-NTD^wt^ from 68.5□ to 77.5□(Fig. 5a), signifying a significant enhancement in thermal stability (ΔTm = 9□). Conversely, ILF marginally changed the thermal stability of the MtbClpC1-NTD by merely 0.5□(Fig. 5a) compared to ILE. The stronger affinity of ILE over ILF correlates with its lower MIC. These findings suggest potential interactions of ILE and ILF with MtbClpC1-NTD. To further test whether these interactions are specific and mutation-dependent, we performed DSF assays using a panel of MtbClpC1-NTD variants harboring the acquired mutations from Mtb resistant strains (F80I/L/V/C). Notably, all four mutations markedly reduced the thermal shift induced by ILE (Fig. 5b - Fig. 5e), indicating that these residues are critical for compound binding. The loss of thermal stabilization strongly supports that resistance arises via disruption of drug-target interaction. Notably, DSF assays of MtbClpX-NTD failed to produce typical sigmoidal melting curves, likely due to the small size and compact nature of the domain (residues 1-112), which may limit the exposure of hydrophobic surfaces upon unfolding. As a result, we could not reliably assess ligand binding to MtbClpX-NTD using this method.

**Fig 5.**
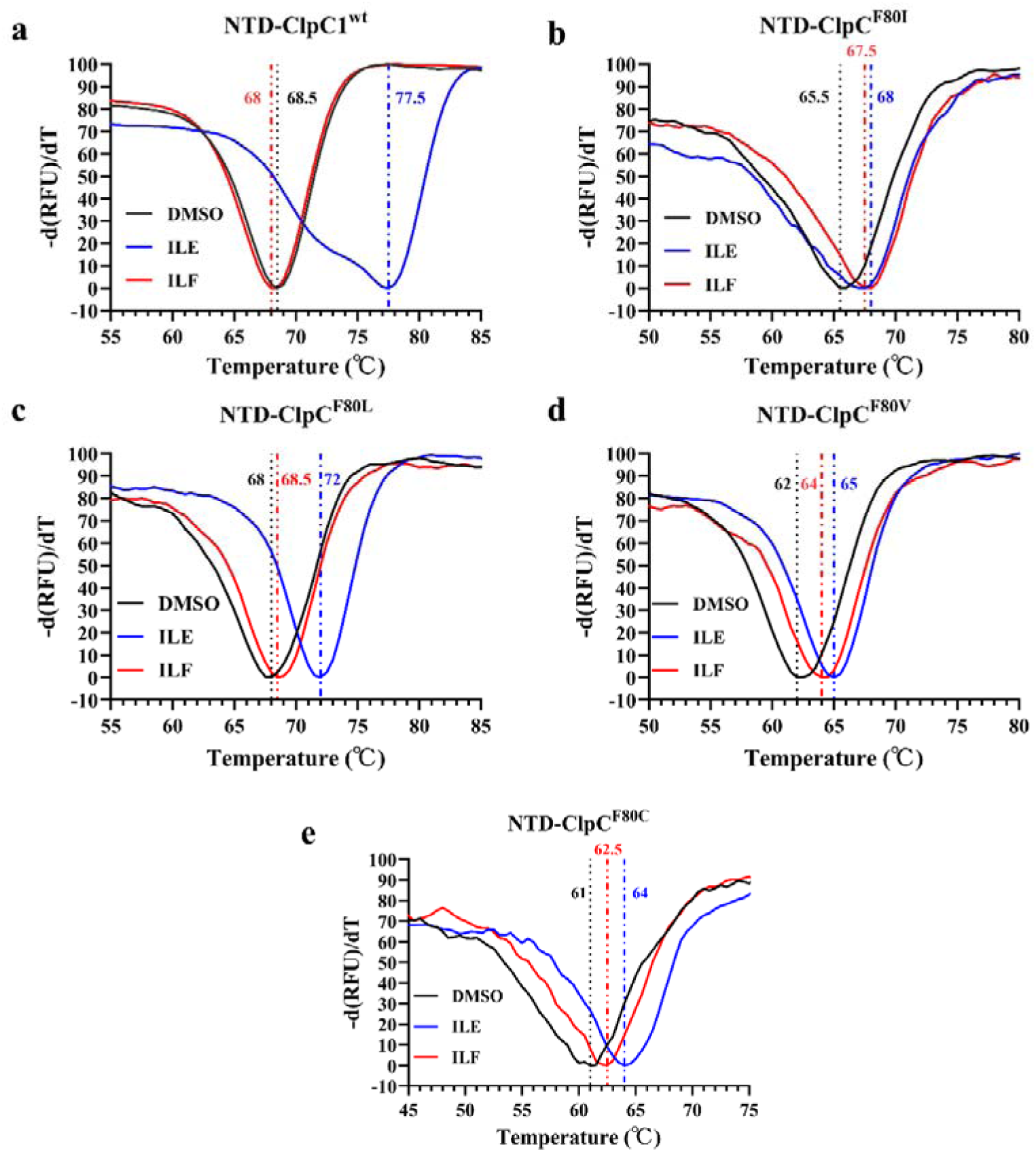
ILE and ILF bind to MtbClpC1-NTD directly. The melting curves of 5 μM MtbClpC1-NTD^wt^ **(a)**, MtbClpC1-NTD^F80I^ **(b)**, MtbClpC1-NTD^F80L^ **(c)**, MtbClpC1-NTD^F80V^ **(d)**, and MtbClpC1-NTD^F80C^ **(e)** in the presence of 10 μM ILE and ILF.

### ILE significantly impedes the proteolytic function of the ClpC1P1P2 and **ClpXP1P2 proteases**

Considering the stronger interactions observed between MtbClpC1-NTD and ILE compared to ILF, we focused our subsequent biochemical assays on evaluating ILE. We observed that ILE inhibited the proteolysis of the model substrate, fluorescein isothiocyanate-casein (FITC-casein), by ClpC1^wt^P1P2 complex compared to the control group (Fig. 6a). However, the proteolytic activities of multiple mutant ClpC1P1P2 proteases remain unaffected at 10 µg mL^-1^ ILE compared to the control, with only partial inhibition noted at 20 µg mL^-1^ ILE (Fig. 6b - Fig. 6e).

**Fig 6.**
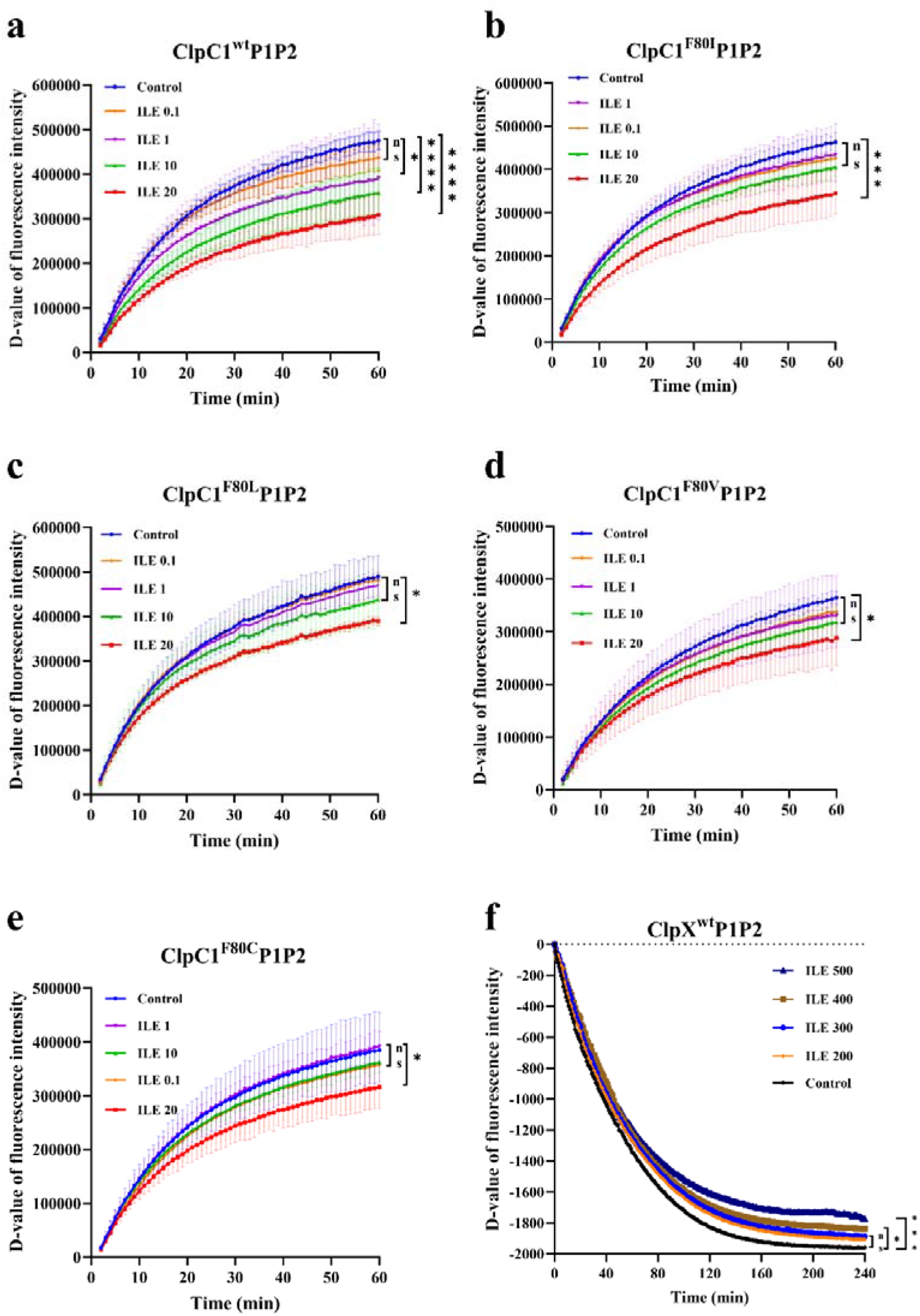
ILE significantly impedes the proteolytic function of both the ClpC1P1P2 proteases and the ClpXP1P2 proteases. Degradation of FITC-casein by **(a)** ClpC1^wt^P1P2, **(b)** ClpC1^F80I^P1P2, **(c)** ClpC1^F80L^P1P2, **(d)** ClpC1^F80V^P1P2, and **(e)** ClpC1^F80C^P1P2 proteases in the presence or absence of ILE. Data are presented as mean ± SD from three biological replicates (n = 3). **f** Degradation of GFP-DAS+4 by ClpX^wt^P1P2 proteases in the presence or absence of ILE. Panel (f) shows data from a representative experiment without error bars. Numbers following the drug names indicate the working concentrations (µg mL^-1^). DMSO was added as control. The D-value of fluorescence intensity represents the change in fluorescence intensity by subtracting the initial fluorescence intensity from the fluorescence intensity of subsequent detections. Statistical significance was determined using one-way ANOVA. *, *P* < 0.05; ***, *P* < 0.0001; ****, *P* < 0.00001; ns, no significant difference, where *P* ≥ 0.05.

For the ClpXP1P2 proteasome, we first used ssrA tagged green florescent protein (GFP-ssrA) as the substrate. The results showed that even when the concentrations of ILE or ILF reached as high as 2000 µg mL^-1^, it had no effect on the proteolytic activity of ClpXP1P2 (Supplementary Fig. 6a and Supplementary Fig. 6b). Previous studies have demonstrated that the adaptor protein SspB from *E. coli* is required for efficient degradation of proteins tagged with a mutated form of the ssrA tag DAS+4 by ClpXP1P2 in both *E. coli* and Msm^44,45^. Therefore, we expressed GFP tagged with DAS+4 (GFP-DAS+4) as the substrate and SspB as the adaptor. Importantly, when using DAS+4-tagged substrates, protein degradation could only be observed in the presence of the adaptor SspB (Supplementary Fig. 6c). It was observed a significant inhibition of the proteolytic activity of the ClpX^wt^P1P2 complex during the hydrolysis of GFP-DAS+4 in the presence of SspB when ILE is added (Fig. 6f). However, we discovered that the mutant ClpX^P30H^P1P2 complex was incapable of degrading GFP- DAS+4 (Supplementary Fig. 6d).

Additionally, we noted that ILE did not impact the ATP-dependent enzymatic activities of ClpC1 and ClpX, as no significant differences in ATPase activities were detected across varying ILE concentrations compared to controls (Supplementary Fig. 7).

### ILE disrupts protein homeostasis with accumulation of ribosomal components and toxin-antitoxin (TA) systems

To elucidate the adaptive responses of Mtb to ILE and gain insight into its mechanism of action, we performed quantitative proteomic analysis of Mtb cells treated with ILE (Supplementary Table 4). Compared to untreated controls, a total of 773 proteins were upregulated and 280 proteins were downregulated (Fig. 7a), indicating a global increase in protein abundance.

**Fig 7.**
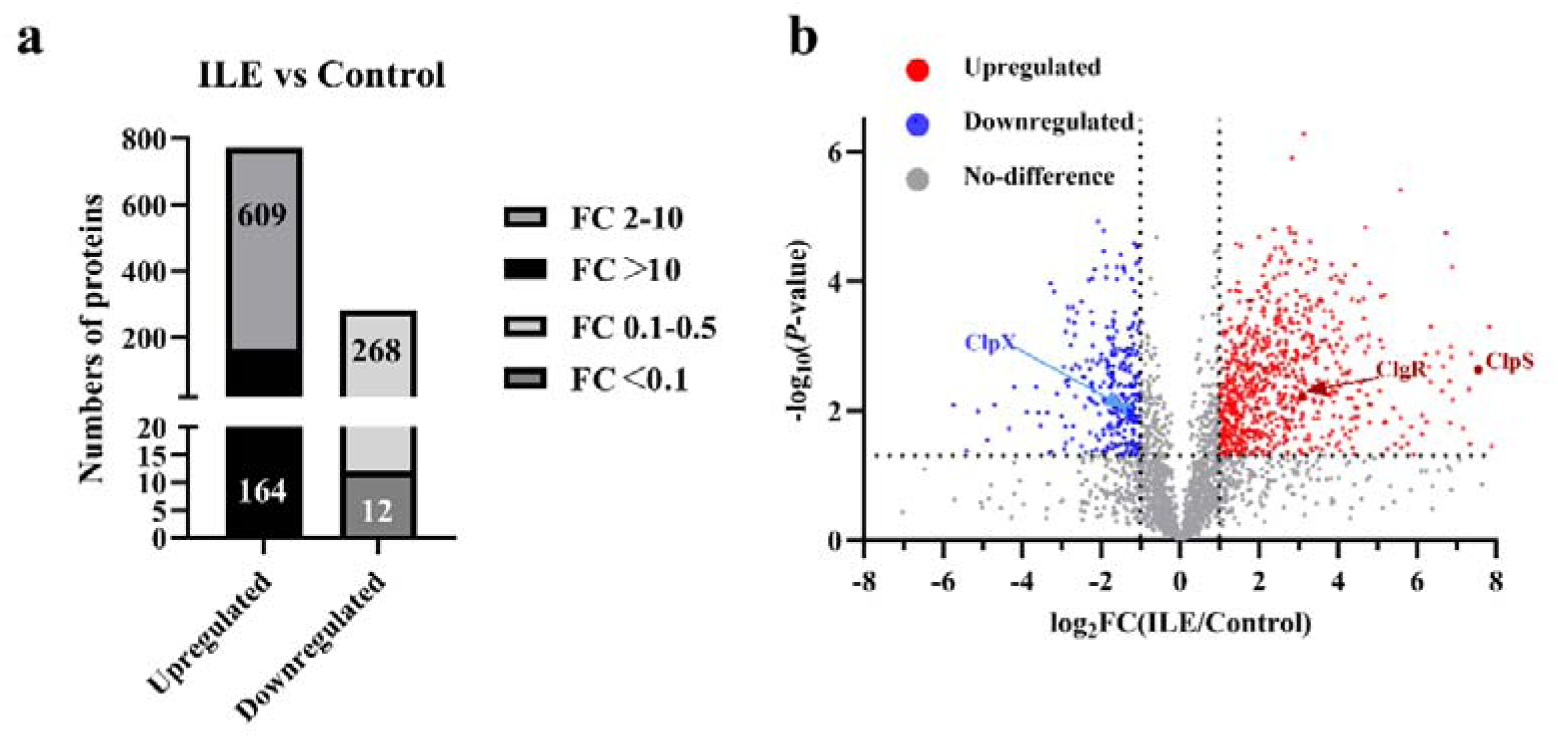
Proteomic profiling reveals global protein expression changes of Mtb in response to ILE treatment. **a** Bar graph summarizing the number of differentially expressed proteins upon ILE treatment compared to the control group. Upregulated and downregulated proteins are categorized by fold change (FC): FC 2-10, FC >10, FC 0.1-0.5, and FC <0.1. **b** Volcano plot showing the distribution of differentially expressed proteins.

Notably, ribosomal proteins—including components of the 50S and 30S subunits (e.g., RpsG, RpsS)—were broadly upregulated, with 21 out of 23 ribosomal proteins showing increased abundance. RplR and RpsC were the only ribosomal proteins found to be downregulated.

All differentially expressed TA system proteins were upregulated, including proteins from VapBC (27 proteins), MazEF (11 proteins), ParDE/RelBE (7 proteins), HigBA (3 proteins) and mt-PemIK (1 protein) families. Additionally, the reported ClpP1P2 substrates PanD (FC = 2.67, *P* < 0.05) and CarD (FC = 5.99, *P* < 0.001) was also significantly upregulated^46,47^. Proteins associated with cell division, such as FtsZ and FtsE, were upregulated approximately two-fold (*P* < 0.001), consistent with scanning electron microscopy observations.

To visualize changes in proteins functionally associated with the Clp protease system, key components including ClpX, ClpS, and ClgR were highlighted in the volcano plot (Fig. 7b). ClpX was downregulated (FC = 0.41, *P* < 0.05), whereas ClpS and ClgR were highly upregulated (FC = 187.26 and 8.53, respectively; both *P* < 0.01). Other components such as ClpC1, ClpC2, ClpP1, and ClpP2 showed no significant changes and were therefore not highlighted.

## Discussion

The study of natural cyclic peptides has emerged as a focal point in the current research on drugs against Mtb. In comparison to the reported cycloheptapeptides, ILE demonstrates superior anti-Mtb activity. Specifically, RUFI inhibited Mtb growth at a MIC of 60 nM by day 14 when tested with the autoluminescent Mtb strain, while ILE achieves an MIC of 48 nM (equivalent to 0.05 µg mL^-1^) by day 15 (Fig. 1a)^48^. Moreover, the MIC of ILE is significantly lower than those of CYMA (100 nM), ECU (160 nM), and lassomycin (410 nM) reported^13,18^. While these compounds exhibit activity against a range of NTM, only RUFI and ILE exhibit efficacy against Mab, indicating their potential as therapeutic agents for diseases caused by Mab^15^.

Previous research had not explored the combination effects of cycloheptapeptides with anti-TB drugs, leaving their potential as adjuvants to current TB therapies unresolved^14–16,18^. Notably, our findings on combinatorial activity suggest that ILE have the potential to enhance the efficacy of existing anti-TB drugs, including RIF, INH, EMB, and TB47. Proteomic analysis provided mechanistic insights into these interactions, revealing that ILE treatment modulated the expression of multiple proteins involved in pathways targeted by these drugs. As for RIF, ILE-induced downregulation of Rho and NusA, two key transcriptional regulators that interact with RNA polymerase. Given that Rho suppression causes widespread transcriptional disruption and rapid cell death, and that NusA modulates transcriptional pausing and termination, their downregulation may sensitize Mtb to RIF-induced RNA polymerase inhibition^49,50^. In the case of INH, four proteins—SigL, Ndh, FadE24, and Glf—were differentially expressed in response to ILE. Specifically, SigL and Ndh were significantly upregulated, whereas FadE24 and Glf were downregulated. SigL has been reported to regulate the expression of KatG, which is crucial for the activation of INH, and Ndh has been identified to influence the NADH/NAD+ ratio, affecting INH-NAD formation^51–53^. INH-resistant Mtb has been found to carry mutations in FadE24^54^. Upregulation of Glf was hypothesized to contribute to INH resistance in Mtb either by binding to a modified form of INH or by sequestering a factor such as NAD+ required for INH activity^55^. For EMB, ILE downregulated UbiA, a key enzyme in the decaprenylphosphoryl-D-arabinose biosynthesis pathway required for cell wall assembly. Given that UbiA overexpression has been associated with EMB resistance, its suppression by ILE may underlie the observed partial synergy^56^. Additionally, synergy with TB47—a cytochrome bc1 complex (QcrB) inhibitor—can be explained by ILE-mediated suppression of several electron transport chain components, including QcrA, QcrC, and CydD^24,57^. As the Qcr and Cyt-bd branches of the electron transport chain are functionally redundant, simultaneous inhibition by TB47 and ILE likely causes a collapse in respiratory function, resulting in pronounced synergistic killing^24^.

Although ILE and ILF have demonstrated potent activity against Mtb, their MOAs remained elusive^17^. Previous studies had identified ClpC1 as the primary target of several cyclic peptides^32^. Consistent with this, we found that mutations in ClpC1— particularly in its NTD—confer resistance to ILE and ILF across multiple mycobacterial species, with the exception of unique insertion mutations discovered between M1 and F2 in both Mab and Msm^15,16,18,36^. Previous studies indirectly validated these mutations via overexpression^36^, but using advanced mycobacterial gene-editing tools, we now directly confirm that specific mutations (e.g., Mab-F2L, Msm-F2C, and a novel Msm insertion) confer ILE/ILF resistance. Additionally, WGS of 2 ILE-resistant Mtb mutants revealed identical large insertions in Rv1572c, a conserved hypothetical protein whose role remains to be explored. Moreover, 3 ILE- resistant Mab mutants were found to carry mutations in *hisS* or *hisD*, which encode histidyl-tRNA synthetase and histidinol dehydrogenase, respectively. Although overexpression of mutated version of these genes did not restore drug sensitivity, gene editing-based validation is planned to confirm their potential role in resistance.

Interestingly, no ClpX mutations had been reported in mutants resistant to these compounds^15,16,18^. However, our findings indicate that ILE-resistant mutants harboring ClpX mutations also exhibit cross-resistance to ILF, suggesting a shared resistance mechanism. Moreover, gene editing confirmed that ClpX mutations can directly confer high-level resistance to both ILE and ILF, revealing ClpX as a novel contributor to cyclic peptide’s resistance. These findings also shed light on possible limitations of previous resistance screening approaches. Whether the ClpX mutation can lead to resistance of mycobacteria to other ClpC1-targeting cyclic peptides—such as RUFI, ECU, and lassomycin—remains to be determined. Given RUFI’s close structural similarity to ILE/ILF and strong activity against Mab, this is particularly worth investigating^15,58^. Molecular docking and gene editing techniques provide effective strategies to explore this possibility.

The MOA of ILE differs from ECU and lassomycin, which stimulate ClpC1 ATPase activity but hinder substrate degradation, as well as CYMA, which activates both ATPase and proteolytic functions^16,18,36^. In this study, we demonstrated that ILE and ILF inhibited the proteolytic activity of both ClpC1P1P2 and ClpXP1P2 proteases without affecting ATPase activity, which is similar to RUFI’s mechanism^15^. Notably, a higher concentration of ILE is required to inhibit ClpXP1P2 than ClpC1P1P2, aligning with the higher resistance levels associated with ClpX mutations. These observations highlight ClpX mutation as a previously unrecognized resistance mechanism in mycobacteria, warranting further investigation.

Proteomic analysis further revealed a broad upregulation of proteins in response to ILE treatment, consistent with a disruption of proteostasis resulting from ClpC1 and ClpX inhibition. Impaired protease activity would prevent normal turnover of these proteins, resulting in their accumulation. This may explain the observed increase in ribosomal protein levels as a compensatory response to maintain translational capacity. Notably, multiple TA systems—including VapBC, MazEF, ParDE/RelBE, HigBA, and mt-PemIK families—were strongly upregulated, aligning with their known degradation by ClpC1P1P2 or ClpXP1P2^9,59^. Given their roles in growth modulation, dormancy, and virulence, such dysregulation could contribute to the antibacterial effects of ILE treatment^60^. Additionally, accumulation of PanD and CarD, two validated ClpP1P2 substrate, further supports protease inhibition^46,47^. ClpX was downregulated, potentially relieving repression of FtsZ and explaining its upregulation^40^. This is consistent with previous reports that ClpX knockdown leads to filamentous cell morphology due to disrupted FtsZ regulation in mycobacteria^41^, and may explain the elongated cells we observed under SEM in ILE/ILF-treated strains. ClpS was dramatically induced, likely to compensate for impaired substrate recognition^59^; and the transcriptional activator ClgR was upregulated, consistent with proteostasis stress responses observed under proteasome inhibition^61^. These findings indicate a coordinated regulatory response aimed at restoring protein homeostasis under Clp protease dysfunction.

However, we discovered that the mutant ClpX^P30H^P1P2 complex was incapable of degrading GFP-DAS+4. This may be attributed to the fact that the enhancement of ClpXP1P2 degradation by SspB requires an intact zinc binding domain (ZBD) in ClpX (residues 1-60)^62^. A mutation in P30, which is located in the hydrophobic surfaces of ZBD, may suppress the binding of SspB to ClpX, thereby preventing the delivery of the tagged protein to ClpXP1P2 complexes for degradation. Although SspB homologs have not been identified in mycobacterium, it is possible that native adaptor-like mechanisms exist, and disruption of such interactions may contribute to ILE resistance. Notably, despite the presence of ClpC1^wt^, the ClpX^P30H^ mutation confers resistance, suggesting that it may compensate for ClpC1 dysfunction by altering substrate specificity or proteostasis regulation. Additionally, the roles of some *clpC1* mutation sites in ILE and ILF resistance remain unproven due to the inability to acquire corresponding gene-edited strains. Therefore, further studies are needed to identify native ClpX substrates or adaptors in mycobacteria, assess ClpX mutant proteolytic activity in more physiological contexts, and attempt to obtain the remaining *clpC1*-edited strains to confirm their roles in ILE and ILF resistance.

While ClpX is not currently considered the target of ilamycins in Mtb, its presence in humans—unlike ClpC1—raises concerns about potential off-target effects. Therefore, we performed sequence analysis shows low similarity (< 25% identity) and no conservation at resistance-related residues between Mtb and human ClpX, suggesting minimal risk. However, further validation is warranted to fully exclude off-target interactions.

In summary, the potent and broad antimycobacterial activities of ILE and ILF underscore their potential for further development as effective agents against mycobacteria, especially drug-resistant strains. Our findings not only reinforce the critical role of the ClpC1P1P2 protease as a validated target, but also uncover ClpX— via mutation—as a previously unrecognized mechanism of resistance^15,16,18,36^. This discovery significantly expands the current understanding of resistance pathways beyond *clpC1*, and suggests that ClpXP1P2 itself represents a promising and underexplored target for antimycobacterial therapy. A dual-targeting strategy that affects both simultaneously is a rational approach, providing a new avenue for developing treatments against mycobacterial infections.

## Materials and Methods

### Bacterial strains and culture conditions

The bacterial strains used in this study are listed in Supplementary Table 5. Mtb, Mab, Mmr, Msm, and derived strains were grown in Middlebrook 7H9 broth (BD, USA) or 7H10 agar (BD) or 7H11 agar (Acmec, China) supplemented with OADC (comprising 0.005% oleic acid, 0.5% bovine serum albumin, 0.2% dextrose, 0.085% catalase). Mtb, Msm, and Mab were grown at 37□. Mmr were grown at 30□. For *E. coli*, *Enterococcus faecium*, *Staphylococcus aureus*, *Acinetobacter baumannii*, *Klebsiella pneumoniae* and *Pseudomonas aeruginosa*, cells were grown in Luria-Bertani (LB) broth at 37□. Antibiotics were supplemented as required at the appropriate screening concentrations for distinct bacterial species. Specifically, for *E. coli* and Msm, the antibiotics added were kanamycin (KAN, 50 µg mL^-1^, Solarbio, China), apramycin (APR, 50 µg mL^-1^, MeilunBio, China), and Zeocin (ZEO, 30 µg mL^-1^, InvivoGen, France). For Mmr, only APR (50 µg mL^-1^) were utilized. Meanwhile, for Mab, KAN was used at a higher concentration of 100 µg mL^-1^, along with ZEO (30 µg mL^-1^) and an increased concentration of APR (230 µg mL^-1^).

### Determination of MICs and MBC against mycobacteria *in vitro* and *ex vivo*

The MICs of ILE and ILF were determined using a cost-effective *in vitro* assay as previously described against a selectable marker-free AlRa, AlRv, AlMab, AlMsm and AlMmr^23–25,30^. The MIC_lux_ was defined as the lowest concentration that could inhibit > 90% relative light units (RLUs) compared with that from the untreated controls^23–25,30^. The MICs of the ILE were also determined by a well-established microplate alamar blue assay (MABA) against Mtb H_37_Ra, H_37_Rv and nine clinical Mtb strains isolated from pulmonary TB patients hospitalized at Guangzhou Chest Hospital, Guangzhou, China^29^. The MIC, as determined by the MABA method, is defined as the lowest concentration that prevents the color change from blue to pink, as observed with the naked eye.

MBC was defined as the minimum concentration resulting in ≥ 99.9% reduction in CFUs compared to the bacterial load before drug addition. For CFU enumeration, mycobacterial cultures treated with different concentrations of compounds or control were collected before or after incubation, with the incubation time varying depending on the species. Bacterial suspensions were serially diluted in phosphate-buffered saline (PBS, GENOM, China), plated onto 7H10 agar plates, and incubated at 37°C until visible colonies appeared. The number of visible colonies was counted, and CFUs were calculated to evaluate bacterial viability.

Intracellular survival assay was performed as previously reported^63^. RAW264.7 macrophages were seeded at 5 × 10□cells per well in 12-well plates and infected with different Mab at an MOI of 10:1 for 4 h. After removing extracellular bacteria with 250 µg mL^-1^ amikacin for 2 h, infected cells were cultured with or without ilamycins as well as 200 ng mL^-1^ aTc. At 72 h post-infection, macrophages were lysed, and intracellular bacteria were quantified by CFU enumeration on 7H10 agar. All experiments were performed in triplicate.

### MICs to non-mycobacteria determination

The MICs of ILE or ILF against *Enterococcus faecium*, *Staphylococcus aureus*, *Acinetobacter baumannii*, *Klebsiella pneumoniae* and *Pseudomonas aeruginosa* were determined by observing the turbidity after 16 h of incubation in LB broth at 37□. The MIC was defined as the lowest concentration, resulting in clarity in turbidity relative to that of untreated control cultures^64–66^.

### Combined activity of ILE or ILF with other compounds *in vitro*

The combined activity of ILE or ILF with other compounds, such as INH (MeilunBio), RIF (MeilunBio), EMB (MeilunBio), STR (MeilunBio), CFZ (MeilunBio), or BDQ (biochempartner, China), and TB47 (BojiMed, China), was evaluated *in vitro* using the checkerboard method. The FICI was interpreted as follows: synergistic (FICI of ≤ 0.5), partially synergistic (0.5 < FICI < 1.0), additive (FICI = 1.0), indifferent (1.0 < FICI ≤ 4.0), and antagonistic (FICI > 4.0)^67,68^.

### Determination of MIC and TC_50_ in macrophage culture

The cell density of mouse macrophages RAW264.7 was adjusted to 5 × 10^5^ cells mL^-1^. 100 μL of cells per well were added to 96-well plates and cultured overnight for adherence. The AlRv concentration was adjusted to 3 × 10^6^ RLU mL^-1^, and 100 μL of AlRv was added to each well. The plates were then incubated at 37□for 6 h. The cells were washed three times with DMEM (Gibco, USA) to remove non-phagocytosed AlRv and treated with DMEM containing amikacin (100□µg mL^-1^, MeilunBio, China) for 2□h to eliminate the extracellular bacteria. The cells were washed twice with DMEM to remove residual amikacin. A total of 195 μL of DMEM containing 10% fetal bovine serum and 5 μL of the two-fold dilutions of various ILE concentrations were added to the cells. The mixture was incubated at 37□with 5% CO_2_, and the RLUs were measured on days 1-5 or 7 of incubation. RIF (1 µg mL^-1^) and dimethyl sulfoxide (DMSO, Xilong Chemical, China) were used as positive and negative controls, respectively.

The TC_50_ of ILE was assessed in mouse RAW264.7 macrophages using the Cell Counting Kit-8 (CCK-8, MeilunBio, China). Cells were seeded in 96-well plates at a density of 1□×□10□cells per well and allowed to adhere overnight. Serial dilutions of ILE were added and incubated for 24□h at 37□with 5% CO_2_. CCK-8 reagent was then added according to the manufacturer’s instructions, and absorbance was measured at 450□nm using a microplate reader. The TC□□ value was calculated from dose–response curves using GraphPad Prism.

### ILE- or ILF-resistant mutants

Broth cultures of AlRa, AlRv, AlMab, and AlMmr, OD_600_ reaching 0.6-1.3, were plated on 7H11 plates containing ILE at 0.1, 0.5, 2.5, 5, 10 or 20 μg mL^-1^. Broth cultures of AlMsm (OD_600_ 0.6-1.3) were plated on 7H11 plates containing ILF at 10, 20 or 40 µg mL^-1^. The bacterial colonies grown on the ILE- or ILF-containing plates were selected and grown in liquid culture for drug susceptibility testing to confirm the drug resistance phenotype.

### WGS analysis

WGS analysis was conducted on the parent strains of Mtb, Mab, and a subset of the confirmed ILE-resistant mutants. The sequencing was performed by the Beijing Genomics Institute in China. Subsequently, the obtained reads were aligned with the reference genome sequence and compared to that of the parent strain. Mutations identified through WGS in the drug-resistant mutants were validated by amplifying and sequencing the *clpX* and *clpC1* genes using Sanger sequencing. Additionally, the *clpX* and *clpC1* genes from other ILE- or ILF-resistant mutants, including Mtb, Mab, Msm, and Mmr, were amplified and sequenced using Sanger sequencing (primer sequences are provided in Supplementary Table 6).

### Molecular docking analysis

To evaluate the binding affinities and interaction modes among ILE, MtbClpC1- NTD, and MtbClpX-NTD, we employed Autodock Vina 1.2.2, a computational protein-ligand docking software previously reported^33^. The structure of ILE was obtained from PubChem Compound^34^. The 3D coordinates of MtbClpC1-NTD were retrieved from the Protein Data Bank (PDB) with PDB code 6CN8 and a resolution of 1.4 Å, while the structure of MtbClpX-NTD was predicted using AlphaFold2.

For the docking analysis, all protein and ligand files were converted to the PDBQT format; water molecules were excluded, and polar hydrogen atoms were supplemented. A grid box was positioned to envelop the domain of each protein, facilitating unrestricted molecular movement. The dimensions of the grid box were set to 30 Å × 30 Å × 30 Å, with a grid point spacing of 0.05 nm. The molecular docking experiments were executed using Autodock Vina 1.2.2, which is accessible at http://autodock.scripps.edu/.

### Overexpression of *clpX* or *clpC1* in Mab, Mmr or Msm

The *clpC1*^wt^, *clpC1*^mt^, *clpX*^wt^ and *clpX*^mt^ from Mtb, Mab, Mmr and Msm were amplified using primers detailed in Supplementary Table 3 and subsequently cloned into the plasmid p60A or pMVA with multi-copy under the control of the *hsp60* promoter as depicted in Supplementary Fig. 3. The plasmids were then transformed into AlMab, AlMmr, and AlMsm. Additionally, the *clpP1*, *clpP2* and *clpX* or *clpC1* of Mab were amplified from the genome of Mab^wt^ and inserted into the plasmid pMVA under the control of the *hsp70* or *hsp60* promoters to enable the simultaneous expression of ClpXP1P2 or ClpC1P1P2. The MICs of the recombinant strains to ILE or ILF were determined using the aforementioned protocols.

### Silencing *clpX* or *clpC1* genes in Mab or Msm

The CRISPR-Cas9 system was employed for targeted gene silencing of *clpX* and *clpC1*^42^. Small guide RNAs (sgRNAs) of 20 base pairs were designed to target the coding regions of *clpX* and *clpC1* on their sense strands. The oligonucleotides were annealed and then ligated with linearized pLJR962^42^. Plasmids harboring sgRNAs against *clpX* or *clpC1*, along with the empty vector pLJR962 as a control, were introduced into Mab and Msm cells via electroporation (Supplementary Table 7).

Tenfold serial dilutions of Mab^wt^ or Msm^wt^, the control strain carrying the empty vector, and the gene-silenced strains were prepared after reaching an OD_600_ of approximately 0.9. For Mab, 1 μL of each dilution was spotted on plates containing either 0.25, 0.5, and 1 µg mL^-1^ ILE or 5, 10, and 20 µg mL^-1^ ILF with varying concentrations of aTc (0, 100, 200 and 400 ng mL^-1^). For Msm, 1 μL of each dilution was spotted on plates containing either 0.004, 0.008, and 0.01625 µg mL^-1^ ILE or 0.125, 0.25, and 0.5 µg mL^-1^ ILF with varying concentrations of aTc (0, 12.5, 25 and 50 ng mL^-1^). Control plates received 1 μL of each dilution with aTc alone. The plates were incubated at 37□ for 3 days. The experiment was performed in triplicate and repeated twice.

### CRISPR/Cpf1-mediated gene editing of *clpX* and *clpC1* genes in Mab and Msm

The CRISPR/Cpf1-associated recombination have been previously employed for gene editing in both Mab and Msm recently^38,39^. Initially, the pJV53-Cpf1 plasmid was electroporated into Mab^wt^ and Msm^wt^ strains, resulting in the Mab::pJV53-Cpf1 and Msm::pJV53-Cpf1, respectively. Subsequently, crRNAs were designed to target specific 25 nucleotide sequences adjacent to the 5’-YTN-3’ motif on the template strand of the target genes (Supplementary Table 7). Oligonucleotides for crRNA expression were ligated into the pCR-zeo vector. These constructed vectors were then electroporated into the respective acetamide-induced cells of Mab::pJV53-Cpf1 or Msm::pJV53-Cpf1. For the induction of Cpf1 expression, aTc was added to 7H11 agar plates to achieve a final concentration of 200 ng mL^-1^. These plates were cultured at 30□ for 3-5 days. Individual colony was picked and screened to confirm successful editing at the target gene site via PCR and Sanger sequencing (primers listed in Supplementary Table 6). The MICs of the edited strains to ILE and ILF were determined using the methods described earlier.

### Expression and purification of proteins

The Mtb ClpP1 (residues 7-200) with a C-terminal Strep-II tag and ClpP2 (residues 13-214) with a C-terminal His_6_-tag were expressed and purified. Briefly, *clpP1* and *clpP2* genes were inserted into the pETDute-1 vector and expressed in *E. coli* BL21 (λDE3) in LB broth at 25C, following induction with 0.5 mM isopropyl β- D-1-thiogalactopyranoside (IPTG, Macklin, China) for approximately 16 h. Cells were resuspended and lysed; The target protein was subsequently purified via nickel affinity chromatography and eluted with the same buffer supplemented with 200 mM imidazole. Eluted fractions containing the target protein were loaded onto a Strep-II column. The target protein was eluted with the strep buffer containing 50 mM HEPES-KOH (pH 7.5), 150 mM KCl, 10 mM MgCl_2_, 50 mM d-Desthiobiotin and 10% glycerol (v/v), before a final gel filtration using on a SuperoseTM 6 increase 10/300 GL column (GE Healthcare) in SEC buffer (25 mM HEPES-KOH (pH 7.5), 150 mM KCl, 10 mM MgCl_2_).

The sequence of *clpX*^wt^ or *clpX*^P30H^ was inserted into the pGEX-6P-1 vector, which had a His_6_-tag at the C-terminal and an HRV 3C protease cleavage site. After induction for 18 h at 16□, the cells were collected by centrifugation. The pellet was then resuspended and lysed. After centrifugation, the supernatant was loaded onto a GST column and eluted. The eluent underwent digestion at 4□ for 26 h with enough HRV 3C protease. The fractions containing the target protein were then pooled, concentrated to 1 mL, and purified with a SuperoseTM 6 increase 10/300 GL column. The sequence of *clpC1*^wt^, its mutants (F80I/V/L/C), the gene encoding *gfp-ssrA* (AADSHQRDYALAA), *gfp*-DAS+4 (AANDENYSENYADAS) or *E. coli sspB* was inserted into the pET-28a vector that had an N-terminal His_6_-tag. The recombinant strains were cultured in LB broth at 37□ until the OD_600_ reached 0.6-0.8. After 18 h of induction with 0.5 mM IPTG at 18□, the pellet was suspended and lysed. Following the loading of the supernatant onto a Ni-NTA column, the proteins were washed and then eluted. The fractions were then pooled and loaded onto a SuperoseTM 6 increase 10/300 GL column.

The gene encoding MtbClpC1-NTD^wt/mt^ or MtbClpX-NTD^wt/mt^ was inserted into the pET-28a vector with a C-terminal His_6_-tag and an HRV 3C protease cleavage site. The expression and purification methods of MtbClpC1-NTD and MtbClpX-NTD were consistent with the methods described above.

### DSF

The binding of ILE or ILF to the MtbClpC1-NTD and MtbClpX-NTD was detected using DSF, following established protocols^43,69^. A 5 µM solution of either MtbClpX^wt/mt^-NTD or MtbClpC1^wt/mt^-NTD was incubated with varying concentrations of ILE or ILF (ranging from 10 to 400 µM), as well as DMSO for 30 minutes, subsequently mixed with Sypro Orange fluorescent dye (Invitrogen, USA). Fluorescence measurements were recorded using the CFX96 Real-Time System (Bio-rad, USA), which involved heating 8-tube PCR Strips from 25□ to 95□ at a constant rate of 0.4□ per minute. The excitation and emission filters were adjusted to wavelengths of 460 nm and 510 nm, respectively. Data collection was facilitated by the CFX Manager software, with analysis performed using GraphPad Prism version 8.3.0. To ensure accuracy, all measurements were performed in triplicate.

### Proteolytic activity assays of ClpC1P1P2 and ClpXP1P2

The proteolytic activities of ClpC1^wt/mt^P1P2 and ClpX^wt/mt^P1P2 were measured at 37□ in black 96-well plates using a Multimode Plate Reader. For the ClpC1P1P2 proteolytic activity experiments, ILE was added at final concentrations of 0, 0.1, 1, 10, and 20 μg mL^-1^ in each well. A 100 µL reaction mixture was formulated with 2 µM ClpP1P2, 1 µM ClpC1^wt/mt^, 2 mM ATP, and 2.5 µM FITC-casein (AAT Bioquest, USA).

For the ClpXP1P2 proteolytic activity experiments, two different ssrA-tagged GFP substrates were used: GFP-ssrA and the mutated ssrA-tagged GFP (GFP-DAS+4). For the GFP-ssrA substrate, the reaction mixture (100 µL) contained 0.2 µM ClpP1P2, 0.7 µM ClpX, 2 mM ATP, 1 µM GFP-ssrA, and 7 µM bortezomib (AbMole, USA) as an activator^70^. For the DAS+4-tagged substrate, GFP-ssrA was replaced with 1 µM GFP-DAS+4, and 2 µM SspB adaptor protein was added.

The hydrolysis of FITC-casein, GFP-ssrA and GFP-DAS+4 was continuously monitored at 535 nm (with excitation at 485 nm). Measurements were performed in triplicate.

### ATPase activity assays of ClpC1 and ClpX

The ATPase activities of Mtb ClpC1 and ClpX were assessed using a free-phosphate detection assay with the Biomol Green reagent (Enzo Life Sciences, USA). To each well of a transparent 96-well plate, 5 μL of various concentrations of ILE (0, 0.08, 0.16, 0.31, 0.625, 1.25, 2.5, 5 and 10 μg mL^-1^) was added. Subsequently, either ClpX or ClpC1 was diluted to a concentration of 0.1 μg/mL in the reaction mixture. 90 μL ClpC1 and ClpX were then added to a 96-well plate and incubated at 37□ for 1 h. The reaction was initiated by adding 5 μL of 1 mM ATP to the reaction mixture, which was maintained at 37□ for 1 h. The final volume of the reaction mixture was adjusted to 100 µL. Post reaction, 100 μL of Biomol Green reagent was introduced to determine the amount of free phosphate. After a 30-minute incubation at room temperature, the resulting reaction product was measured at 620 nm using a Multimode Plate Reader. The concentration of free phosphate liberated from ATP by ClpX and ClpC1 was calculated using a standard curve based on the known concentration of free phosphate. Measurements were performed in triplicate.

### Microscopy of drug-treated Msm

Treatment of exponentially growing Msm was conducted using 0.125 μg mL^-1^ ILE, 5 μg mL^-1^ L ILF, or 1 μg mL^-1^ L CLR (Meilunbio, China) for a duration of 15 h. The bacteria were immobilized with a 4% glutaraldehyde solution, followed by three washes and resuspension in PBS. Bacterial suspensions were then applied to 5 mm cell slides, which were placed in a 24-well plate to dry naturally. Afterward, the cell slides underwent three additional PBS washes. The cell slides were subsequently

soaked in ascending concentrations of ethanol solutions (70%, 80%, 90%, 100%) for 10 minutes each. Finally, the cell slides were dried using the critical point drying method and coated with an electrically-conducting material. Observations of bacterial morphology were made using a GeminiSEM 300 ultra-high resolution field emission cryo-scanning electron microscope.

### Proteomic analysis

The AlRa cells were treated with 0.0125 μg mL^-1^ of ILE for 5 days in triplicate for 4D label-free phosphoproteomic analysis. To maintain comparable cell density between the untreated and ILE-treated groups, the untreated cells were diluted on the same day as the compound addition, prior to sample collection. The cells were resuspended and boiled for 20 minutes. Subsequently, the samples were sent to APTBIO (Shanghai, China) for proteomic testing and analysis.

To compare the abundance of phosphopeptides between the control and treatment samples, label-free quantification was performed with a minimum FC of 2 to determine the differentially expressed phosphopeptides. Additionally, the student’s *t*-test was employed to detect significant differences between the control and treatment samples, with *P* < 0.05 indicating significance. rotein functional annotation was conducted using NCBI and Mycobrowser (https://mycobrowser.epfl.ch/). All differentially expressed proteins were listed in Supplementary Table 4.

### Data availability

The Sequence Read Archive (SRA) for a subset of ILE-resistant Mab strains have been deposited in the National Center for Biotechnology Information (NCBI) under BioProject Name PRJNA1262689. For the remaining samples, only variant analysis results provided by the sequencing company are available and are included in Supplementary Table 2 and 3. All other data are available from the corresponding author on reasonable request.

## Acknowledgments

We acknowledge the team of professor Yicheng Sun from the Institute of Pathogenic Biology, Chinese Academy of Medical Sciences for kindly sending the pJV53-Cpf1 and pCR-Zeo plasmids as the tool of gene editing. This work was supported by fundings from National Key R&D Program of China (2021YFA1300900), National Natural Science Foundation of China (81973372, 32300152, 82022067, 22037006), China Postdoctoral Science Foundation (2022M723164), Guangdong Provincial Basic and Applied Basic Research Fund (2022A1515110505, 2019B030302004), Grant of State Key Lab of Respiratory Disease (SKLRD-Z-202414, SKLRD-Z-202301), Open Program of Shen Zhen Bay Laboratory (SZBL2021080601006) and Nansha District Science and Technology Plan Project (NSJL202102).The funders had no role in study design, data collection and analysis, decision to publish, or preparation of the manuscript.

## Author contributions

T.Z., J.J., and X.X. supervised the project; T.Z., Y.G., and C.F. designed the whole project and most of the experiments; J.M. and C.S. prepared and characterized the ilamycin E and ilamycin F; Y.G. and C.F. performed the susceptibility testing and resistant mutant screening; B.Z. and J.J. contributed to molecular docking simulation; C.F., X.T., and J.H. constructed overexpression and gene-editing strains and peformed the susceptibility testing; C.F. and X.H. performed the gene-scilencing experiments; Y.G. and B.Z. contributed to protein expression and mode of action experiments; C.F. and H.Z. contributed to the microscopy results; Y.G. performed the statistical analysis; Y.G., F.C., and T.Z. wrote the manuscript; H.M.A., J.L., J.M., X.C., N.Z., X.X., J.J., and T.Z. revised the manuscript.

