## Supplementary Fig,1~7, Supplementary Table 3,5,6,7 for "Mutations in ClpC1 or ClpX subunit of the caseinolytic protease confer resistance to natural product ilamycins in mycobacteria"

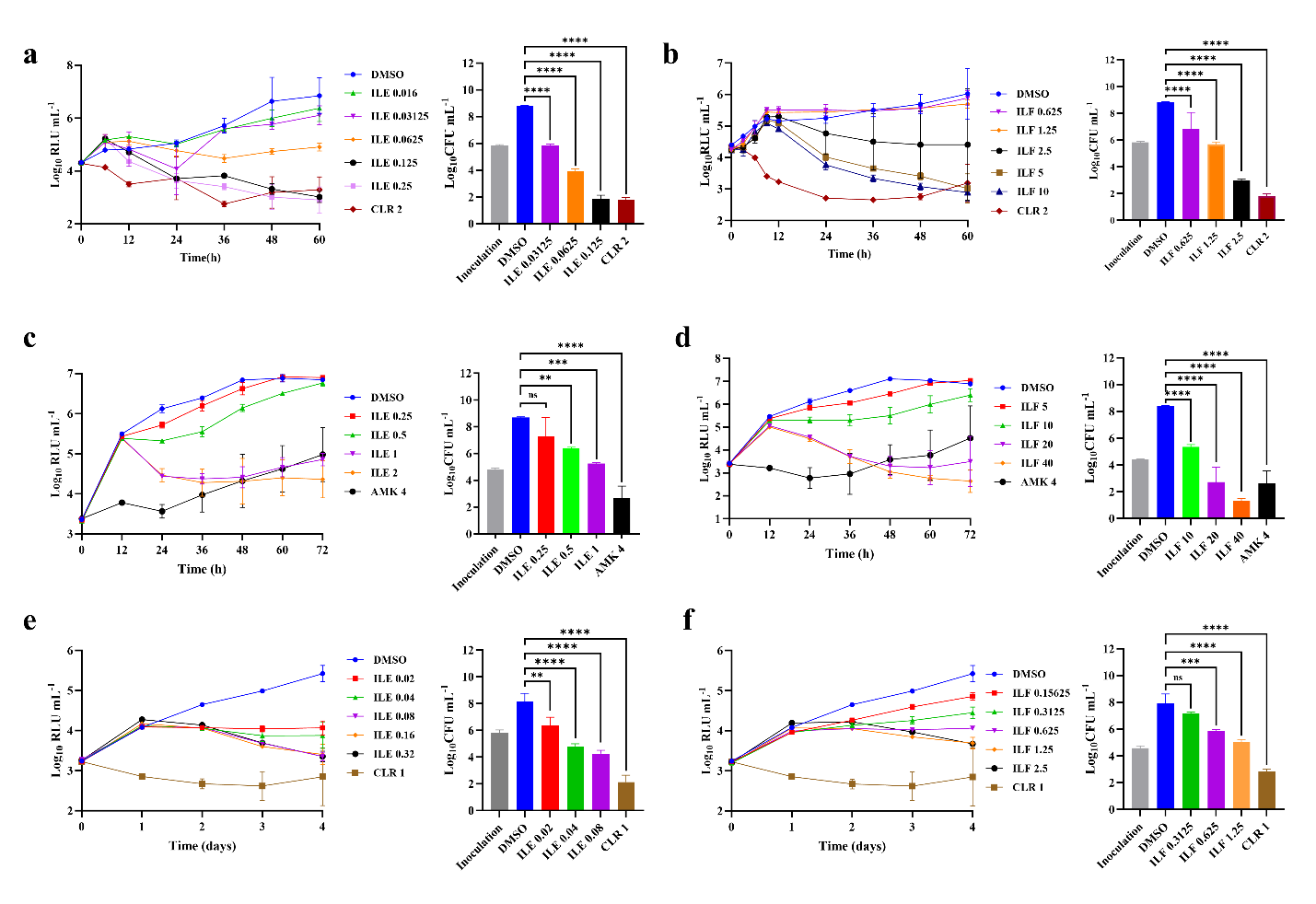

**Supplementary Fig 1** **| Time-kill curves and CFU counts of ILE and ILF against autoluminescent nontuberculous mycobacteria.** Time-kill curves and CFU counts of ILE against AlMsm **(a)**, ILF against AlMsm **(b)**, ILE against AlMab **(c)**, ILF against AlMab **(d)**, ILE against AlMmr **(e)**, and ILF against AlMmr **(f)**. DMSO, dimethyl sulfoxide, solvent control; CLR, clarithromycin, positive control for AlMsm and AlMmr; AMK, amikacin, positive control for AlMab. All drug concentrations are in μg mL^-1^. The MIC_lux_ was defined as the lowest concentration that can inhibit > 90% relative light units (RLUs) compared with that from the untreated controls. RLU and CFU data were obtained from independent experiments. Statistical significance was determined using one-way ANOVA. Data are presented as mean ± SD from three biological replicates (n = 3), ns, not significant; ^*^, *P* < 0.05; ^**^, *P* < 0.01; ^***^, *P* < 0.001; ^****^, *P* < 0.0001.

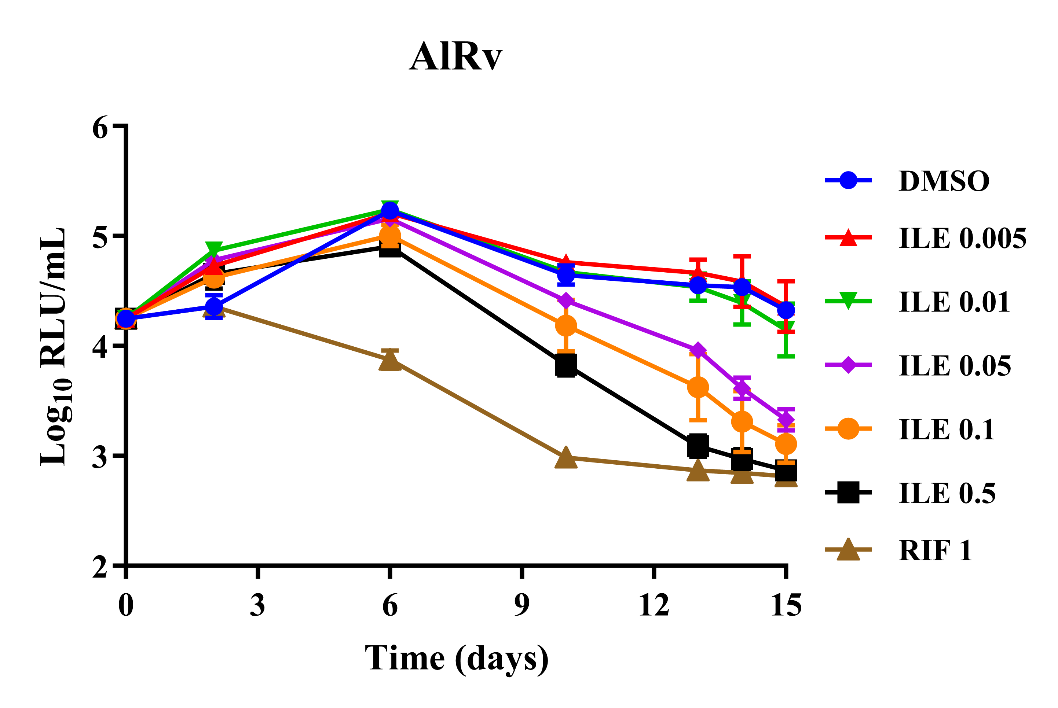

**Supplementary Fig 2 | Time-kill kinetics of ILE against AlRv.** DMSO, solvent control; RIF, positive control. All drug concentrations are in μg mL-1. The MIClux was defined as the lowest concentration that can inhibit > 90% relative light units (RLUs) compared with that from the untreated controls. Data are presented as mean ± SD from three biological replicates (n = 3).

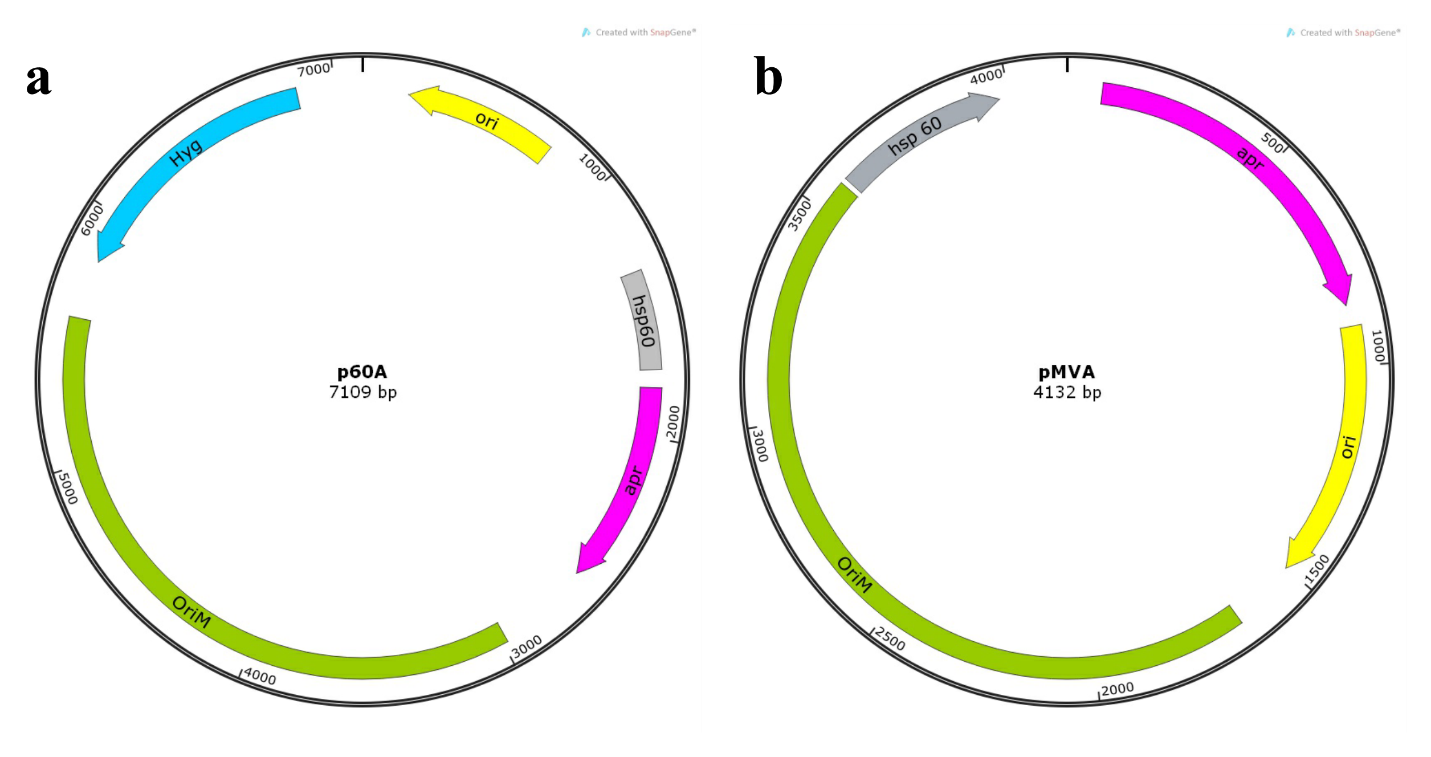

**Supplementary Fig 3 | Maps of overexpression vectors with different resistance marker.** **a** p60A. **b** pMVA. *hsp60*, a mycobacterial strong promoter; *Ori*, high-copy-number origin of replication in *E. coli*; *OriM*, origin of replication in Mycobacteria; *apr*, aparamycin resistant gene; *hyg*, hygromycin B resistant gene.

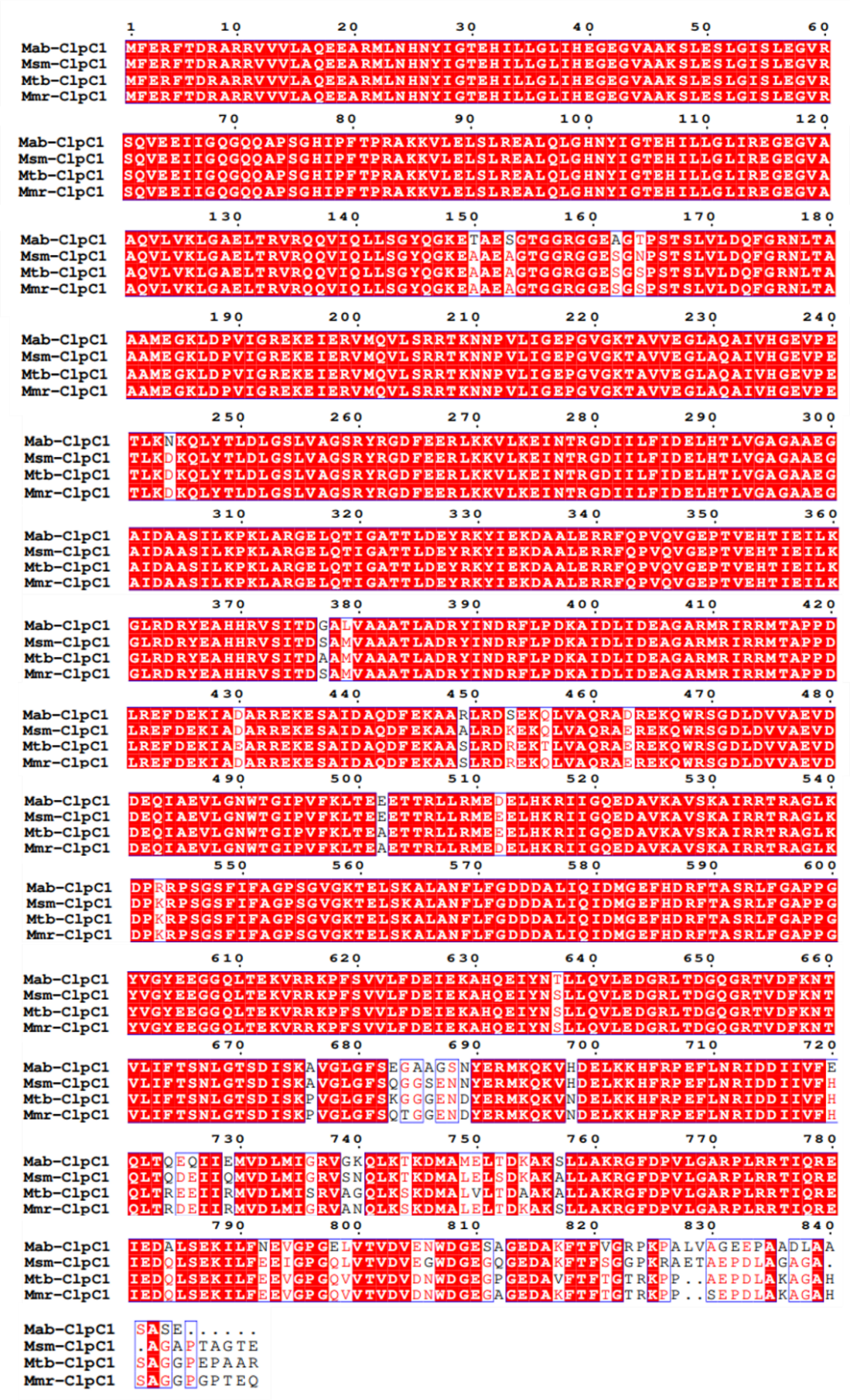

Supplementary Fig 4 | Alignments of amino acid sequences of ClpC1 across Mtb, Mab, Mmr, and Msm

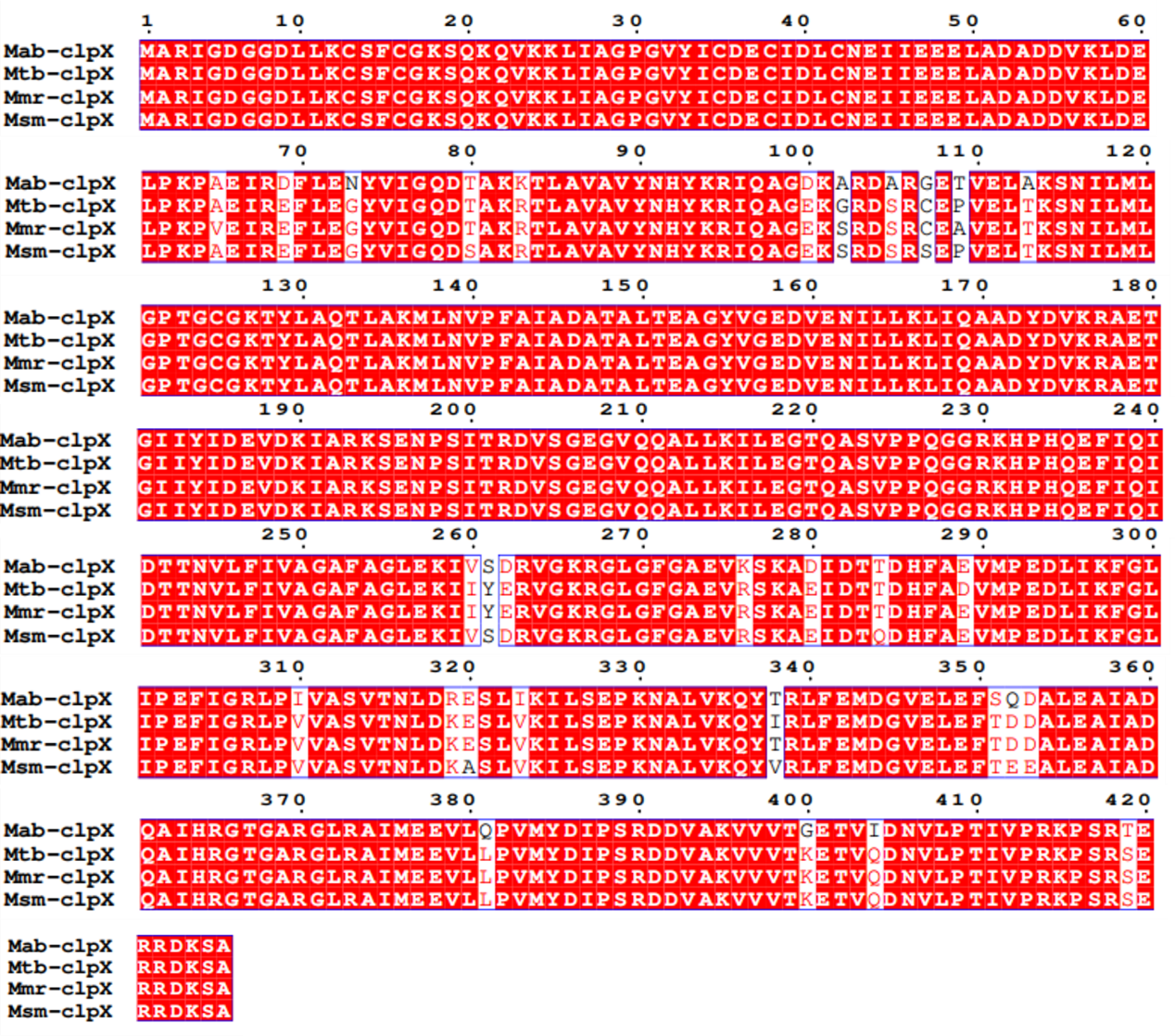

Supplementary Fig 5 | Alignments of amino acid sequences of ClpX across Mtb, Mab, Mmr, and Msm

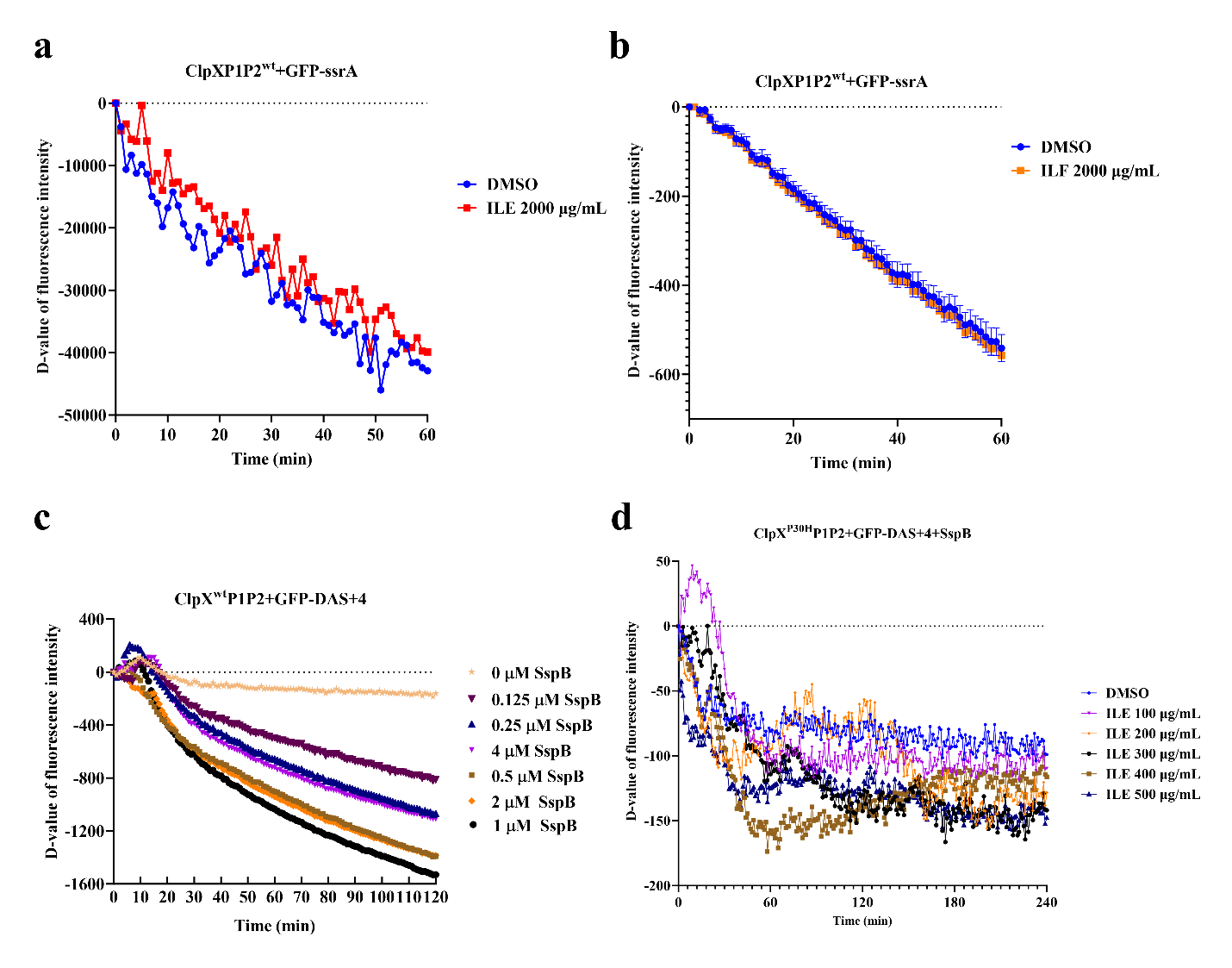

**Supplementary Fig 6 | Effects of ILE and ILF on ClpXP1P2 protease activity using GFP-tagged substrates.** Time-course degradation curves of GFP-ssrA by ClpX^wt^P1P2 in the presence of 2000 μg mL⁻¹ ILE (**a**) or ILF (**b**). **c** Degradation of GFP-DAS+4 by ClpX^wt^P1P2 in the presence of increasing concentrations of the adaptor SspB (0 to 4 μM), confirming that degradation of DAS+4-tagged substrates requires the presence of SspB. **d** Inhibition of GFP-DAS+4 degradation by ClpX^P30H^P1P2 in the presence of 2 μM SspB under increasing concentrations of ILE (0–500 μg μg mL^-1^). Data are shown as Δ-values of fluorescence intensity over time.

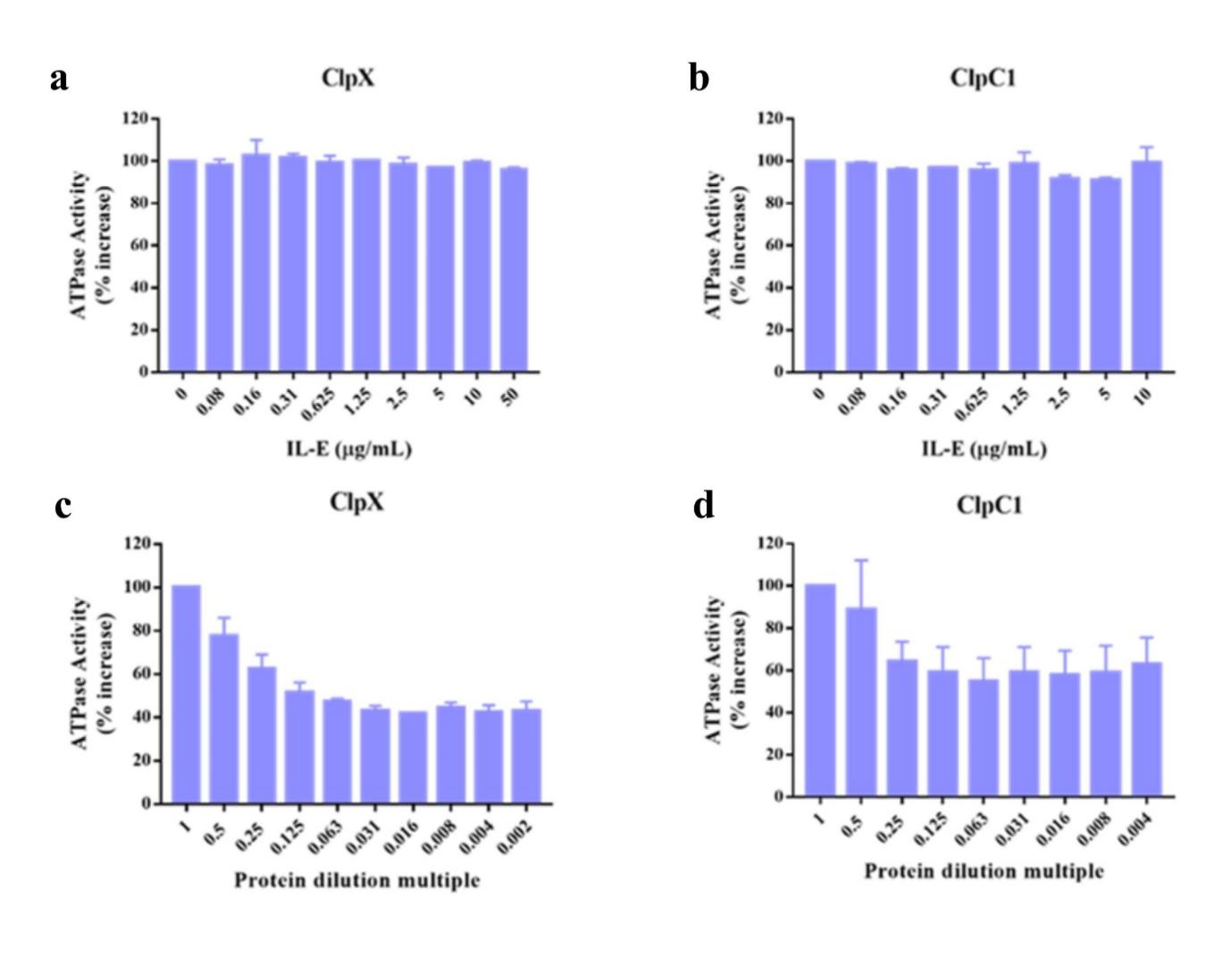

**Supplementary Fig 7** **| Effect of ILE on the ATPase activities of ClpX and ClpC1. a** Effect of ILE on the ATP-dependent ATPase activities of ClpX. **b** Effect of ILE on the ATP-dependent ATPase activities of ClpC1. Data are mean ± SD, *n*=3.

**Supplementary Table 3 | MICs of ILE and ILF against ClpC1 and ClpX overexpressing strains**

| **Strains** | **MICs (μg/mL) / (fold change)** | |
| --- | --- | --- |
|  | **ILE** | **ILF** |
| AlMmr^wt^ | 0.08 | 0.625 |
| AlMmr::ClpC1^wt^ | 0.08 / (1) | 0.625 / (1)  -  F80L  TTT→CTT  -  -  F80C  TTT→TGT  -  -  -  -  P30H  CCC→CAC  F2L  TTC→TCC  -  - |
| AlMmr::ClpC1^L88R^ | 0.625 / (8) | 5 / (8)  -  F80L  TTT→CTT  -  -  F80C  TTT→TGT  -  -  -  -  P30H  CCC→CAC  F2L  TTC→TCC  -  - |
| AlMmr::ClpC1^F2L^ | 0.08 / (1) | 0.625 / (1) |
| AlMmr::ClpC1^H77D^ | 0.08 / (1) | 0.625 / (1) |
| AlMab^wt^ | 1 | 20 |
| AlMab::ClpX^wt^ | 1 / (1) | 20 / (1) |
| AlMab::ClpX^P30H^ | 1 / (1) | 20 / (1) |

**Supplementary Table 5 | Bacterial strains used in this study**

| **Strains** | **Description** | **Source or reference** |
| --- | --- | --- |
| AlRa | Selectable marker-free autoluminescent Mtb H_37_Ra | ^1^ |
| AlRv | Selectable marker-free autoluminescent Mtb H_37_Rv | ^2^ |
| AlMsm | Selectable marker-free autoluminescent Msm mc^2^155 | ^1^ |
| AlMab | Selectable marker-free autoluminescent Mab | ^3^ |
| AlMmr | Selectable marker-free autoluminescent Mmr | ^4^ |
| H_37_Rv | Widely used virulent laboratory Mtb strain, ATCC 27294 | Lab stock |
| K11 | Drug-resistant Mtb clinical isolate | Guangzhou Chest Hospital, Guangzhou, China |
| K12 | Drug-resistant Mtb clinical isolate | Guangzhou Chest Hospital, Guangzhou, China |
| K13 | Drug-resistant Mtb clinical isolate | Guangzhou Chest Hospital, Guangzhou, China |
| K14 | Drug-resistant Mtb clinical isolate | Guangzhou Chest Hospital, Guangzhou, China |
| K16 | Drug-resistant Mtb clinical isolate | Guangzhou Chest Hospital, Guangzhou, China |
| K17 | Drug-resistant Mtb clinical isolate | Guangzhou Chest Hospital, Guangzhou, China |
| K18 | Drug-resistant Mtb clinical isolate | Guangzhou Chest Hospital, Guangzhou, China |
| K19 | Drug-resistant Mtb clinical isolate | Guangzhou Chest Hospital, Guangzhou, China |
| K20 | Drug-resistant Mtb clinical isolate | Guangzhou Chest Hospital, Guangzhou, China |
| K11 | Drug-resistant Mtb clinical isolate | Guangzhou Chest Hospital, Guangzhou, China |
| *Staphylococcus aureus* | Common Gram-positive pathogen, clinical isolate | Guangzhou Chest Hospital, Guangzhou, China |
| *Enterococcus faecium* | Common Gram-positive pathogen, clinical isolate | Guangzhou Chest Hospital, Guangzhou, China |
| *Acinetobacter baumannii* | Common Gram-negative pathogen, clinical isolate | ^5^ |
| *Klebsiella pneumoniae* | Common Gram-negative pathogen, clinical isolate | ^6^ |
| *Pseudomonas aeruginosa* | Common Gram-negative pathogen, clinical isolate | ^7^ |
| Mab^wt^ | Mab GZ002 | ^8^ |
| Msm^wt^ | Msm mc^2^155 | Lab stock. |
| Mab::pJV53-Cpf1 | Mab GZ002 containing plasmid pJV53-Cpf1 | ^9^ |
| Mab-C1^F2L^ | Mab GZ002 in which c*lpC1*^wt^ was edited into c*lpC1*^F2L^ | This study |
| Mab-X^P30H^ | Mab GZ002 in which c*lpX*^wt^ was edited into c*lpX*^P30H^ | This study |
| Msm::pJV53-Cpf1 | Msm mc^2^155 containing plasmid pJV53-Cpf1 | ^10^ |
| Msm-C1^F2C^ | Msm mc^2^155 in which c*lpC1*^wt^ was edited into c*lpC1* ^F2C^ | This study |
| Msm-C1^M1-L-F2^ | Msm mc^2^155 in which c*lpC1*^wt^ was edited into c*lpC1* ^M1-L-F2^ | This study |
| Msm-C1^I197N^ | Msm mc^2^155 in which c*lpC1*^wt^ was edited into c*lpC1*^I197N^ | This study |

Supplementary Table 6 | Primers used in this study for PCR

| **Primers** | **Primer sequences (5’-3’)** | **Purpose** |
| --- | --- | --- |
| TB-X-U1 | gccactgtagggtggtcag | To amplify *clpX* gene from ILE-resistant Mtb mutants for checking mutation. |
| TB-X-D1 | gatcagatcctccggcatc |  |
| TB-X-U2 | ggccaagatgcttaacgtgc |  |
| TB-X-D2 | gccatgacgaacccgatcg |  |
| TB-C1-U1 | ggcgacctgacatttggctac | To amplify *clpC1* gene from ILE- resistant Mtb mutants for checking mutation. |
| TB-C1-D1 | cagttcaccgcgagcgagcttc |  |
| TB-C1-U2 | gtcggcaagaccgcggtc |  |
| TB-C1-D2 | gagaacggcttgcgccgc |  |
| TB-C1-U3 | gcggatcatcgggcaagaggac |  |
| TB-C1-D3 | gccgccgatacgtcatctg |  |
| Mab-X-U1 | ggagtaccgcaagctctc | To amplify *clpX* gene from ILE-resistant Mab mutants and gene-edited strains for checking mutation. |
| Mab-X-D1 | gcatgacctctgcgaagtg |  |
| Mab-X-U2 | gccgactacgacgtgaagcg |  |
| Mab-X-D2 | ggccaattcggcaacgcg |  |
| Mab-C1-U1 | gcggccaactagtacgttc | To amplify *clpC1* gene from ILE-resistant Mab mutants for checking mutation. |
| Mab-C1-D1 | gcaccgatggtctgcagctc |  |
| Mab-C1-U2 | ggcgaggtgcccgagacac |  |
| Mab-C1-D2 | gaccgtcctcgagaacctg |  |
| Mab-C1-U3 | gtccaaggcgctggccaacttc |  |
| Mab-C1-D3 | gcactcgtcgaagggccac |  |
| Mm-C1-F1 | cgggtgtatctgattcgggt | To amplify *clpC1* gene from ILE-resistant Mmr mutants for checking mutation. |
| Mm-C1-R1 | gctcgtcgatgaacaggatg |  |
| Mm-C1-F2L-F | gcaagaccgctgtggtcga |  |
| Mm-Cl-R2 | cgaagagcaccacgctga |  |
| Mm-C1-F3 | ccttccggtgtcggtaag |  |
| Mm-C1-R3 | ctcgcctcctcggcacatg |  |
| Mm-X-F1 | cgtacagcgcgtctcggc | To amplify *clpX* gene from ILE-resistant Mmr mutants for checking mutation. |
| Mm-X-R1 | ccgggagccgaccgatga |  |
| Mm-X-F2 | cagcaggccctgttgaag |  |
| Mm-X-R2 | ggcttggtcccagttgttg |  |
| Ms-C1-F | caacttagcggaaggttgcca | To amplify *clpC1* gene from ILF-resistant Msm mutants for checking mutation. |
| Ms-C1-R | cgtgacggacagccacagc |  |
| Ms-X-F | atcggcgcagcacatgagg | To amplify *clpX* gene from ILF-resistant Msm mutants for checking mutation. |
| Ms-X-R | gaccaacggcggttacgg |  |
| Mab-X-CZ-F | caagacaattgccatatgatggcacgtatcggagac | To amplify the *clpX*^wt^ or *clpX*^P30H^ from Mab mutants for cloning into the overexpression vector p60A. |
| Mab-X-CZ-R | ggtcgacggtatcgataagcttctacgcggacttgtc |  |
| Mab-C1-CZ-F | tggccaagacaattgccatatgatgttcgagagattca | To amplify the wt or mutated *clpC1* from Mab mutants for cloning into the overexpression vector p60A. |
| Mab-C1-F2L-CZ-F | tggccaagacaattgccatatgatgtccgagagattca |  |
| Mab-C1-CZ-R | tcgacggtatcgataagcttttactccgaggcggag |  |
| Mm-C1-CZ-F | agacaattgcggatccatgttcgaacgttttaccgaccg | To amplify the wt or mutated *clpC1* from Mmr mutants for cloning into the overexpression vector pMVA. |
| Mm-C1-F2L-CZ-F | agacaattgcggatccatgtccgaacgttttaccgaccg |  |
| Mm-C1-CZ-R | cgacatcgataagcttttactgctcggtgggccc |  |
| Ms-C1-CZ-F | ccaagacaattgcggatccatgtttgagagatttaccgaccgc | To amplify the wt or mutated *clpC1* from Msm mutants for cloning into the overexpression vector pMVA. |
| Ms-C1-F2S-CZ-F | ccaagacaattgcggatccatgtgtgagagatttaccgaccgc |  |
| Ms-C1-2insL-CZ-F | ccaagacaattgcggatccatgtgttttgagagatttaccgac |  |
| Ms-C1-F5I-CZ-F | ccaagacaattgcggatccatgtttgagagaattaccgaccgc |  |
| Ms-C1-CZ-R | cgtcgacatcgataagctttcactccgtgcctgcgg |  |
| hsp70-F | tgcctgcaggtcgactctagagacccgcacgaccagcgt | To amplify the *hsp70* promoter from Mab for cloning into the overexpression vector pMVA. |
| hsp70-R | aacgcatggtgaatcctcctgaatatgtagagc |  |
| Mab-P12-F | aggaggattcaccatgcgttcggcgacagcc | To amplify *clpP1* and *clpP2* genes from Mab for cloning into the overexpression vector pMVA. |
| Mab-P12-R | ggtacccggggatcctctagactacgaagctgtctgagcgga |  |
| Mab-P12-XF | cgggatccatggcacgtatcggagac | To amplify *clpP1* and *clpP2* genes from Mab for cloning into the overexpression vector pMVA. |
| Mab-P12-XR | aactgcagctacgcggacttgtcgc |  |
| Mtb-ClpC1-F | ggaattccatatgatgttcgaacgatttacc | To amplify *clpC1* genes from Mtb for cloning into the overexpression vector p60A. |
| Mtb-ClpC1-R | cccaagcttctaccgcgcggccggctcc |  |
| Mtb-ClpX-F | ggaattccatatgatggcgcgcataggag | To amplify *clpX* genes from Mtb for cloning into the overexpression vector p60A. |
| Mtb-ClpX-R | cccaagcttctacgcgctcttgtcgcg |  |
| X-60-F | aagttctgttccaggggcccgagctgcccaagccg | To amplify *clpX* or mutated clpX genes from Mtb for cloning into the protein expression vector pGEX-6P-1. |
| X-60-R | gggaattccggggatcccagctacgcgctcttgtcgc |  |
| pGEX-PX-F | ctgggatccccggaattccc | To amplify the vector pGEX-6P-1 gene |
| pGEX-PX-R | gggcccctggaacagaactt |  |
| ClpP2-F | aactttaagaaggagatataccatggcgcgctacatc | To amplify Mtb *clpP2* (Remove 30 bases from the N-terminal) gene for cloning into the protein expression vector pETDuet-1. |
| ClpP2-linker-R | ttagtggtgatgatggtgatgggagccaccgccaccagagccaccaccgccggcggtttgcgcggagagctt |  |
| pET-Dute1-F1 | catcaccatcatcaccactaa | The vector pETDuet-1 was amplified in order to recombine with the *clpP2* gene for pETDuet-1-ClpP2 |
| pET-dute1-R1 | ggtatatctccttcttaaagttaaacaaaa |  |
| clpP1-F | taccctcgagtctggtatgcgttcgaactcgca | To amplify Mtb *clpP1* (Remove 18 bases from the N-terminal) gene for cloning into the protein expression vector pETDuet-1. |
| clpP1-R | cgcagcagcggtttctttttacttttcgaactgcgggtggctccactgtgcttctccattgacg |  |
| pET-Dute1-F2 | aaagaaaccgctgctgcg | The vector pETDuet-1-ClpP2 was amplified in order to recombine with the *clpP1* gene for pETDuet-1-ClpP1P2 |
| pET-Dute1-R2 | accagactcgagggtacc |  |
| 28a-CX-F | ctcgagcaccaccaccaccaccactgagatccggctgctaac | The vector pET-28a was amplified in order to recombine with the wt or mutated *clpC1* gene for pET28a-ClpC1. |
| 28a-CX-R | tacaggttctcgtgatgatgatgatgatgcatggtatatctccttct |  |
| 28a-C1-F1 | tcacgagaacctgtactttcaggggatgttcgaacgatttaccgaccgt | To amplify the wt *clpC1* or mutated *clpC1* genes from Mtb for cloning into the protein expression vector pET28a. |
| 28a-C1-F2 | tcacgagaacctgtactttcaggggatgctcgaacgatttaccgaccgt |  |
| 28a-C1-R2 | gtggtggtgctcgaggccgatggacctagaccg |  |
| 28a-3C-F | gtttcagggcccgcaccaccaccaccaccactgagatccggct | The vector pET28a was amplified in order to recombine with the Mtb *clpC1* (1-435bp) or Mtb *clpX* (1-336bp) gene for pET28a-ClpC1-NTD or pET28a-ClpX-NTD. |
| 28a-3C-R | ccttcttaaagttaaacaaaattatttctagaggg |  |
| TBC1-3CN-F | gtttaactttaagaaggagatataccatgttcgaacgatttaccgacc | To amplify Mtb *clpC1* (1-435 bp) gene for cloning into the protein expression vector pET-28a. |
| TBC1-3CN-R | gtggtgcgggccctgaaacagcacttccaggtaaccggagagcagctggatc |  |
| TBX-3CN-F | gtttaactttaagaaggagatataccatggcgcgcataggagac | To amplify Mtb *clpX* (1-336 bp) gene for cloning into the protein expression vector pET-28a. |
| TBX-3CN-R | gtgcgggccctgaaacagcacttccagcaactcaacgggctcacatcg |  |
| PET-GFP-CZ-R | aatgggtcgcggatccgtgagcaagggcgag | To amplify *gfp* gene for cloning into the protein expression vector pET-28a. |
| GFP-ssrA-CZ-R | tcgagtgcggccgcaagcttttaagctgctaaagcgtaatcacgttg |  |
| GFP-DAS+4-CZ-R | gtgcggccgcaagctttgaggcatctgcataattttccgaatagttctcatcgttcgccgccttgtacagctcgtccat |  |
| SspB-CZ-F | aatgggtcgcggatccatggatttgtcacagctaacaccac | To amplify *sspB* gene from *E. coli* for cloning into the protein expression vector pET-28a. |
| SspB-CZ-R | gtgcggccgcaagcttttacttcacaacgcgtaatgccg |  |

Supplementary Table 7 | The crRNAs, sgRNAs and oligonucleotides used for gene editing and gene silencing.

| **crRNAs, sgRNAs and oligonucleotides (5’-3’)** | **Description** |
| --- | --- |
| gagagattcactgaccgcgccaggc | crRNA targeting the F2 of ClpC1 in Mab |
| ccgcctggcgcggtcagtgaatctctcgaGcatcggtctccctcgcttcctcgtccggc | oligonucleotides targeting the lagging strand of *clpC1* in Mab for editing into *clpC1*^F2L^ |
| atcgccggccccggcgtgtacatct | crRNA targeting the P30 of ClpX in Mab |
| cgatgcattcatcgcagatgtacacgccgTggccggcgatgagcttcttgacctgcttc | oligonucleotides targeting the lagging strand of *clpX* in Mab for editing into *clpX*^P30H^ |
| agagatttaccgaccgcgcc | crRNA targeting the F2 of ClpC1 in Msm |
| ccctgcgggcgcggtcggtaaatctctcaCacatcggtggttacctgcttccatcacgt | oligonucleotides targeting the lagging strand of *clpC1* in Msm for editing into *clpC1*^F2C^ |
| cctgcgggcgcggtcggtaaatctttcGaaTAAcatcggtggttacctgcttccatcac | oligonucleotides targeting the lagging strand of *clpC1* in Msm for inserting M1-L-F2 |
| gatttccttctcgcggccga | crRNA targeting the I197 of ClpC1 in Msm |
| cgatccggtcatcggccgcgagaaggaaaAcgagcgggtcatgcaggtgctgagccggc | oligonucleotides targeting the lagging strand of *clpC1* in Msm for editing into *clpC1*^I197N^ |
| ggtgccgatgtagttgtggt | sgRNA for silencing *clpC1* in Mab |
| tggccgatgacgtagttctc | sgRNA for silencing *clpX* in Mab |
| ggtgccgatgtagttgtggt | sgRNA for silencing *clpC1* in Msm |
| tgaccgatgacgtaaccctc | sgRNA for silencing *clpX* in Msm |
